# Molecular basis for the recognition of steroidogenic acute regulatory protein by the 14-3-3 protein family

**DOI:** 10.1101/2020.04.14.040436

**Authors:** Kristina V. Tugaeva, James Titterington, Dmitriy V. Sotnikov, Eugene G. Maksimov, Alfred A. Antson, Nikolai N. Sluchanko

## Abstract

Steroidogenesis in adrenals and gonads starts from cholesterol transport to mitochondria by the steroidogenic acute regulatory protein STARD1, containing a mitochondrial import sequence followed by a cholesterol-binding START domain. Although mutations in this protein have been linked to lipoid congenital adrenal hyperplasia, the mechanism of steroidogenesis regulation by the STARD1 remains debatable, hypothetically involving a molten-globule structural transition and interaction with 14-3-3 proteins. We show that, while the isolated START domain does not interact with 14-3-3, interaction is enabled by STARD1 phosphorylation at Ser57, close to the mitochondrial peptide cleavage site. Biochemical analysis of the STARD1 affinity towards 14-3-3 and crystal structures of 14-3-3 complexes with Ser57 and Ser195 phosphopeptides, suggest distinct roles of site-specific phosphorylations in recruiting 14-3-3, to modulate STARD1 activity, processing and import to mitochondria. Phosphorylation at Ser195 creates a unique conditional site, that could only bind to 14-3-3 upon partial unfolding of the START domain.

## INTRODUCTION

Steroid hormones control many processes including metabolism, immune functions and sex differentiation. Steroidogenesis in adrenals and gonads is kinetically limited by cholesterol transport to the inner mitochondrial membrane (IMM), where the enzymatic conversion of cholesterol into pregnenolone takes place [1]. In the acute phase, this bottleneck stage of steroidogenesis is largely dependent on the cAMP-dependent *de novo* synthesis and function of the steroidogenic acute regulatory protein StAR, also referred to as STARD1 [2-5]. Reduction, deletion or knockout of the *star* gene arrests steroidogenesis leading to the accumulation of cholesterol-enriched lipid droplets in the cytoplasm of steroidogenic cells [6, 7]. Likewise, a range of STARD1 mutations impair its function leading to autosomal recessive disorders called lipoid congenital adrenal hyperplasias (LCAH) [6]. The critical role of STARD1 in steroidogenesis is underlined by the ability of the wild-type STARD1 transgene to completely rescue the lethal STARD1^-/-^ phenotype in knockout mice [8, 9].

Although STARD1 was discovered as early as in 1983 [4] (identified in 1994 [3]), its complete structure is still unknown and its mechanism of action remains the subject of much debate [1, 10, 11]. STARD1 is produced in our bodies as a pre-protein of 285 amino acids containing an N-terminal mitochondrial targeting sequence (first ∼60 residues) and the so-called steroidogenic acute regulatory lipid-transfer (START) domain able to bind cholesterol (Fig. 1A). The START domain has the α/β helix-grip fold (∼210 amino acids) featuring a lipid-binding hydrophobic cavity [12] (Fig. 1B). This domain is found in 15 members of the START protein family, being present either as a sole domain or as part of a multidomain protein [10, 12], and is responsible for binding various lipids.

**Fig. 1.**
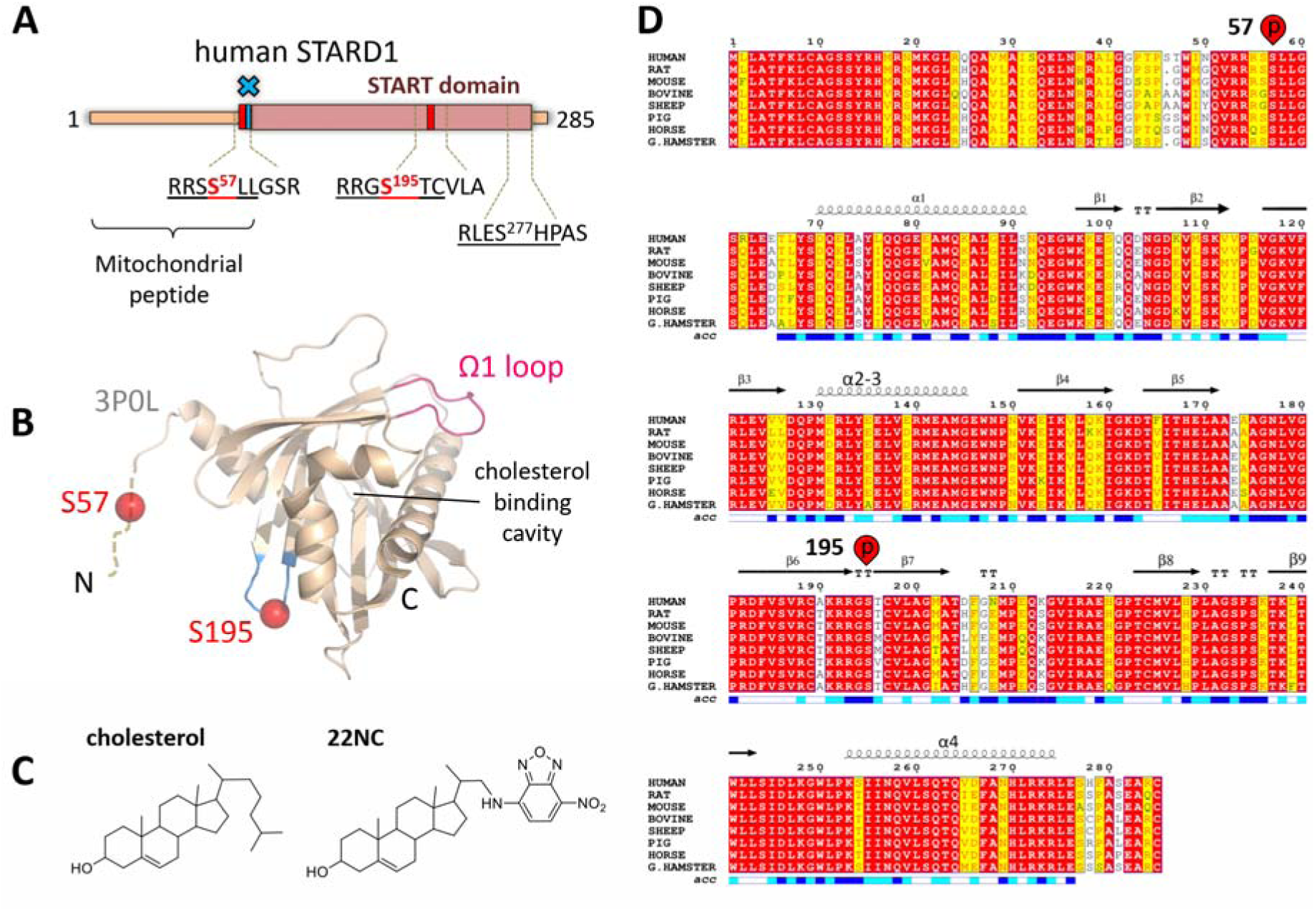
Structure and sequence conservation of STARD1. A. Schematic of the human STARD1. Sequences of the three predicted 14-3-3-binding segments are shown, with central interacting motifs underlined. PKA-phosphorylated sites [32] are in red. Blue cross depicts region of pre-protein processing upon mitochondrial import. B. Ribbon diagram of STARD1 (PDB code 3P0L) showing location of phosphorylation sites (red spheres), cholesterol-binding cavity and Ω1 loop implicated in lipid binding. Note the location of the Ser195 site within the β6-β7 loop (shown in light blue). Ser57 is located within the disordered N-terminal region not resolved in the crystal structure (dashed ribbon). C. Chemical structures of cholesterol and its fluorescent analog 22NC bearing the solvatochromic NBD group. D. Sequence alignment of eight mammalian STARD1 orthologs drawn using ESPript 3.0 [49] showing sequence similarity, secondary structure elements as in (B) and the Ser57 and Ser195 phosphorylation sites. The scale below reflects solvent accessibility (acc) as low (white), intermediate (cyan) or high (blue), calculated by ESPript 3.0 [49] based on the 3P0L crystal structure.

Crystal structures have been determined only for the apo forms of human STARD1 (3P0L) [13], STARD3 (5I9J [14] and 1EM2 [15]) and several other START domains [13]. No START domain structure in complex with cholesterol is available to date, however, flexibility of the Ω_1_ loop and unfolding of the C-terminal α_4_ helix have been implicated in cholesterol binding and release [16-18]. Although the molecular mechanism of cholesterol transfer remained unexplained, several hypotheses were put forward. Initially, it has been proposed that STARD1 transports cholesterol as a shuttle [15]. According to an alternative model of cholesterol transport, STARD1 adopts a molten globule-like conformation in the vicinity of mitochondrial membranes or at a lower pH inside the intermembrane space [19, 20]. A similar concept, involving unfolding of the START domain, has been adopted also for the functioning of STARD6 [21] and STARD3 proteins [22, 23]. Noteworthily, STARD3 is thought to serve as the regulator of steroidogenesis in placenta, the only steroidogenic tissue where STARD1 is not expressed [11, 22]. Cholesterol was suggested to foster a structural transition of STARD1, in a concentration-dependent manner [24, 25]. However, even though STARD1 unfolding was widely observed [19, 23, 26], and was directly linked to STARD1 functionality, it remains unclear how this process can be reversed to ensure delivery of ∼400 cholesterol molecules per min by each STARD1 molecule during the acute phase of steroidogenesis [27]. Moreover, mutational analysis indicated that the ability of STARD1 to bind cholesterol, commonly assessed *in vitro* using fluorescent cholesterol analogs, such as 22-NBD-cholesterol (22NC) (Fig. 1C), was not directly linked to its steroidogenic activity [28].

The N-terminal leader sequence apparently directs the STARD1-mediated cholesterol transfer to mitochondria [29]. The processing of the human STARD1 pre-protein [27, 30, 31], regulated by cAMP concentration [27] and STARD1 phosphorylation [32], is associated with its import to mitochondria, where eventually a processed protein is accumulated [19, 30]. The sequence of these molecular events is unclear due to the absence of a direct correlation between the STARD1 cleavage and steroidogenic activity. Indeed, being immobilized in the outer mitochondrial membrane (OMM), the processed protein devoid of the leader peptide is still capable of promoting steroidogenesis, at least in certain cell types and at specific conditions [31]. This led to the attractive ‘pause-transfer’ hypothesis that for carrying out its function, STARD1 needs to reside in the OMM, until being imported into mitochondria for processing and inactivation [27, 33]. While partially rescuing STARD1^-/-^ phenotype in knockout mice, STARD1 devoid of the first 47 residues failed to provide sufficient cholesterol transport to the mitochondria of steroidogenesis cells *in vivo*, suggesting a more complex role of the mitochondrial targeting peptide in STARD1 functioning [8].

Originally described as a rapidly induced phosphoprotein [4, 5, 34, 35], STARD1 can undergo Ser/Thr phosphorylation [32, 36, 37], which is likely achieved with the help of the A-kinase anchoring proteins and the regulatory subunits of cAMP-dependent protein kinase (PKA) for more effective, location-specific STARD1 phosphorylation at the OMM [38, 39]. Phosphorylation of highly conserved Ser57, located in the N-terminal part of the protein preceding the START domain (Fig. 1A, 1D), apparently had no direct effect on pregnenolone synthesis [32] but may interfere with the STARD1 processing due to Ser57 location close to the cleavage sites [30]. Phosphorylation by PKA at another conserved residue, Ser195 (Fig. 1D), enhanced steroidogenesis by 50% [32]. In contrast to the phosphorylatable wild-type transgene, STARD1 with the S195A amino acid substitution failed to rescue STARD1^-/-^ phenotype in knockout mice, characterized by intracellular accumulation of cholesterol droplets, low level of steroid hormones and neonatal lethality [9]. These effects of Ser195 phosphorylation lacked mechanistic explanation, as modification of Ser195 didn’t change STARD1 folding, cholesterol-binding ability [28, 40] or the mitochondrial import efficiency *in vitro* [32]. The functional role of STARD1 phosphorylation remains largely unclear but likely involves the regulation of STARD1 interaction with other proteins.

The 14-3-3 family proteins have been recently described as direct STARD1 partners and regulators of steroidogenesis [41, 42]. These eukaryotic protein hubs are involved in a variety of regulatory functions via selective binding to protein partners phosphorylated at specific Ser/Thr residues [43], Fig. 2A. 14-3-3 binding can regulate protein activity, subcellular localization or mediate protein-protein interactions [44]. Human 14-3-3 proteins are expressed as 7 different isoforms (β, ζ, ε, σ, τ, γ, η) encoded by different genes [43]. They assemble into homo- and heterodimers [45] and can themselves undergo phosphorylation. Phosphorylation of a semi-conserved serine (Ser58 in 14-3-3ζ, Ser59 in 14-3-3γ), located at the dimer interface, destabilized 14-3-3 dimer and affected protein functionality [46, 47]. Recent data suggested that phosphorylation of this serine in 14-3-3γ as well as 14-3-3 monomerization can affect 14-3-3/STARD1 interaction [41, 42]. Taken together, the available evidence suggest that, depending on STARD1 phosphorylation, 14-3-3 proteins can detain STARD1 in the cytoplasm, thereby regulating steroidogenesis [41, 42]. Yet, the structural basis for the 14-3-3/STARD1 interaction remained unknown.

**Fig. 2.**
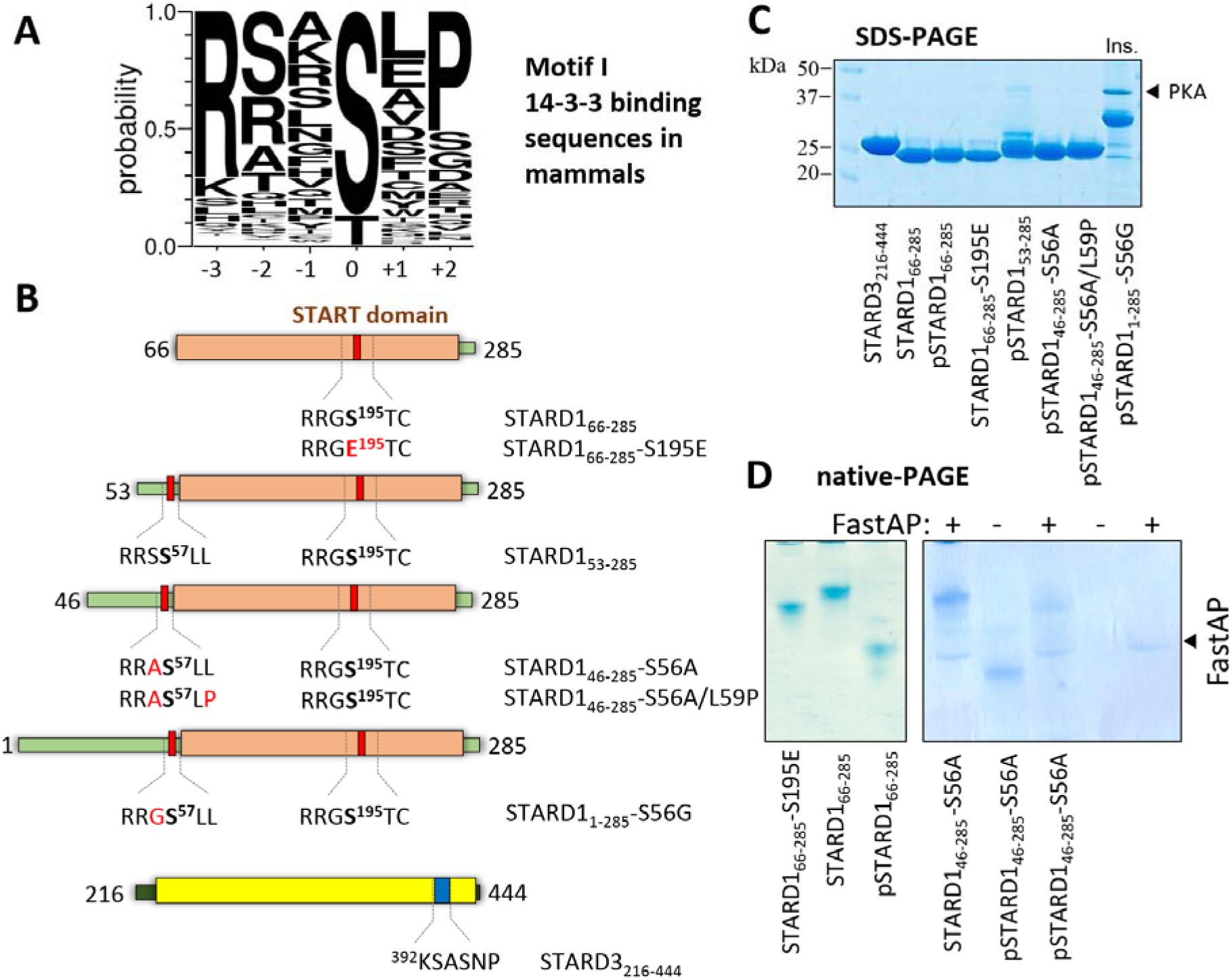
Human STARD1 and STARD3 proteins used in this study. A. Normalized Weblogo [55] diagram showing the consensus 14-3-3 binding sequence motif derived by aligning 217 sequences of mammalian 14-3-3 partners [56]. C-terminal binding motifs were not included. The central residue (position 0) is phosphorylated. B. Scheme showing STARD1 and STARD3 constructs used. The central phosphorylatable STARD1 residues are shown in bold and mutated amino acids in red. C. SDS-PAGE of protein samples used in this work. Note that all constructs were soluble except for the full-length STARD1 (Ins.). D. Native PAGE showing the effect of phosphorylation of several STARD1 constructs. To reverse phosphorylation, phosphatase (FastAP) was added in the “+” samples (FastAP position on the gel is indicated by arrow).

Here, we dissected the binary interaction between 14-3-3 and STARD1 by considering the effects of the oligomeric state of 14-3-3 and phosphorylation of both proteins. In contrast to previous reports [42, 48], we show that START domains of STARD1 and STARD3 in their unphosphorylated, natively folded state, do not bind to 14-3-3, regardless of its oligomeric state. Possessing two conserved PKA phosphorylation sites within sequence motifs suboptimal for 14-3-3-binding, phosphorylated STARD1 itself has low affinity towards 14-3-3, indicating that these two proteins may instead interact as part of a larger protein assembly that forms *in vivo*. Furthermore, crystallographic analysis of 14-3-3σ complexes with the STARD1 phosphopeptides containing the central Ser57 or Ser195 revealed that, while both phosphopeptides bind canonically to 14-3-3, conformations of the Ser195 peptide observed for the complex with 14-3-3 are completely different to its conformation in the unbound STARD1 (PDB 3P0L) where this segment forms a β6-β7 loop [13]. The data point to a unique, conditional binding of STARD1 to 14-3-3 and reveal the remarkable structural polymorphism of STARD1, which may be critical for its interaction with 14-3-3 and, therefore, for the regulation of cholesterol transport and steroidogenesis.

## RESULTS

### Potential 14-3-3-binding sites in human STARD1

STARD1 was first identified as phosphoprotein upgerulated in the acute phase of steroidogenesis in cAMP-stimulated adrenal cortex, corpus luteum and Leydig cells [4, 5, 34, 35]. Phosphorylation of certain Ser/Thr residues is a known prerequisite for 14-3-3 binding [43, 50]. SCANSITE 4.0 predicts that human STARD1 protein (residues 1-285) contains three candidate 14-3-3-binding motifs, namely, RRSS^57^LL, RRGS^195^TC, and RLES^277^HP. The first motif is located within the N-terminal part of the protein, upstream of the START domain, whereas the second and the third motifs reside inside the START domain (Fig. 1A). Out of the three, the RLES^277^HP sequence matches the canonical 14-3-3-binding motifs best (Fig. 2A). However, it is located within C-terminal α-helical sequence that is not conserved among STARD1 orthologs (Fig. 1D). PhosphoSitePlus (www.phosphosite.org) defines Ser277 as a minor phosphorylation site, detected only in one study [51]. In contrary, the RRSS^57^LL and RRGS^195^TC motifs are defined as good targets for PKA [31, 32, 52, 53] and, therefore, may serve as 14-3-3-binding sites in human STARD1.

We constructed several human STARD1 variants (Fig. 2B): i) a standalone STARD1 domain (residues 66-285), ii) a processed STARD1 lacking N-terminal peptide but containing both Ser57 and Ser195 (two constructs, starting from residue 53 or residue 46), and iii) a full-length STARD1 (residues 1-285). Aiming to avoid previously reported problems with STARD1 aggregation [15], we prepared STARD1 variants as fusions with a cleavable maltose-binding protein (MBP) [26]. The proteins were highly pure (Fig. 2C) and showed a significant phosphorylation-related shift on native-PAGE when co-expressed with PKA (Fig. 2D). Importantly, further incubation of phosphoproteins with PKA *in vitro* did not increase electrophoretic mobility whereas addition of phosphatase returned the protein band to the initial position (Fig. 2D), suggesting stoichiometric phosphorylation of the obtained preparations. As expected, the phosphomimicking mutant S195E displayed a smaller shift due to partial charge imitation, simultaneously confirming that Ser195 is efficiently phosphorylated within the intact STARD1 domain (Fig. 2D) [32, 52, 54].

Despite a number of trials that included different expression strains, temperatures, levels of induction, purification in the presence of detergents and protein refolding from inclusion bodies, it proved impossible to obtain soluble full-length STARD1_1-285_. Hence, the longest viable constructs examined in this study lacked the N-terminal residues 1-45.

### Interaction of STARD1 constructs with 14-3-3

First, we questioned whether the START domain can interact with 14-3-3 (Suppementary Fig. 1). As we showed previously, in solution STARD1_66-285_ predominantly exists in a monomeric conformation equivalent to the crystallographic monomer (PDB 3P0L) [54] and is capable of binding steroids [26, 40]. Likewise, we produced and characterized the START domain of the closest STARD1 homolog, STARD3, which, in the context of non-expressed STARD1, is responsible for steroidogenesis in placenta [11, 22]. According to the synchrotron SAXS data, our STARD3 sample revealed a very similar solution structure to that of STARD1 (Supplementary Fig. S2). The START domain of either protein was mixed with 14-3-3, but even the highest protein concentrations did not allow us to detect the interaction using analytical size-exclusion chromatography (SEC). No interaction was observed regardless of whether STARD1 was PKA phosphorylated at Ser195 or not, and regardless of whether the dimeric 14-3-3 or the engineered monomeric 14-3-3 form, m58E mutant [57], were used (Supplementary Fig. 1). A similar result was obtained for the phosphomimicking S195E mutant of STARD1, or a different 14-3-3 isoform (Supplementary Fig. 1). The inability to detect even weak complexes of 14-3-3 with STARD1 phosphorylated at Ser195 was unexpected. Although the structure of the Ser195 containing motif, a short loop connecting β-strands, is inconsistent with the flexible extended conformation required for binding to 14-3-3, it has been proposed to be the main determinant for binding [41].

Likewise, although STARD3 was recently reported to bind 14-3-3 in a phosphorylation-*independent* manner using the 392-KSASNP-397 motif [48] located in the β8-β9 loop (PDB code 5I9J [14]), our sample of STARD3 did not display any interaction neither with the 14-3-3 dimer nor the engineered 14-3-3 monomer (m58E) (Supplementary Fig. 2). Together, these data disfavored the possibility of tight binding of natively folded isolated START domains of STARD1 and STARD3 to 14-3-3 proteins.

Therefore, we extended the STARD1 construct and produced STARD1_53-285_ protein which contained phosphorylatable Ser57, in addition to Ser195. The protein was co-expressed with PKA and produced in a rather low yield. Mass-spectrometry analysis revealed phosphorylation of human STARD1_53-285_ at not only conservative Ser195 and Ser57, but also the neighboring, semi-conservative residue, Ser56 (Supplementary Table 1 and Supplementary Fig. 3). Phosphorylation of these neighboring serines in rat STARD1 was also achieved by PKG [51]. As observed previously for the 14-3-3 interaction with Cdc25B phosphatase [58], such tandem phosphorylation can preclude binding to 14-3-3.

**Table 1.** Quantitation by surface plasmon resonance of the interaction between doubly phosphorylated STARD1_46-285_-S56A/L59P and the His-tagged immobilized 14-3-3.

| 14-3-3 type | $K_D$ (M) | $K_A$ (1/M) | $k_{off}$ (1/s) | $k_{on}$ (1/(M*s)) |
| --- | --- | --- | --- | --- |
| His-tagged 14-3-3ε homodimer | $1.3 \cdot 10^{-5}$ | $7.7 \cdot 10^4$ | $5.8 \cdot 10^{-3}$ | $0.4 \cdot 10^{-3}$ |
| His-tagged 14-3-3$\epsilon/\gamma$ heterodimer | $1.5 \cdot 10^{-5}$ | $6.7 \cdot 10^4$ | $5.7 \cdot 10^{-3}$ | $0.4 \cdot 10^{-3}$ |

Therefore, to avoid the Ser56 phosphorylation detected in STARD1_53-285_ and taking benefit from the presence of alternative residues in close orthologs (Fig. 1D), we have replaced Ser56 with Ala. To improve the yield and solubility, we extended the constructs at the N terminus to residue 46. In addition to the authentic version of the protein, STARD1_46-285_-S56A, we also created construct STARD1_46-285_-S56A/L59P which contained the canonical Pro in the +2 position of the phosphoserine-57 peptide owing to the L59P mutation (Fig. 2). Both versions were co-expressed with PKA and purified to homogeneity similar to STARD1_66-285_ (Fig. 2C, D). Phosphorylation of Ser57 and Ser195 was confirmed by mass-spectrometry (Supplementary Table 1 and Supplementary Fig. 3) and led to a complete shift on native PAGE (Fig. 2D) confirming stoichiometric phosphorylation. This is in line with earlier observations that Ser57 and Ser195 are good substrates of PKA [32, 52] and that its co-expression pushes phosphorylation of the cognate sites to completion [59, 60].

Analytical SEC revealed interaction of 14-3-3 with extended STARD1 constructs phosphorylated at both, Ser57 and Ser195 positions (“pSTARD1”). Interaction of the STARD1_46-285_-S56A/L59P protein was significantly enhanced compared to the STARD1_46-285_-S56A (Fig. 3A). Remarkably, the latter protein construct, representing the authentic processed STARD1, showed interaction with 14-3-3ε and 14-3-3γ isoforms that are up-regulated in the acute phase of steroidogenesis [41], but complexes were insufficiently stable to give a distinct peak on the elution profile (Fig. 3B). In contrast, titration of 14-3-3 by phosphorylated STARD1_46-285_-S56A/L59P ensured the appearance of a peak corresponding to the protein complex and allowed rough estimation of its apparent Mw as ∼93 kDa (Fig. 3C), indicating 2:2 binding stoichiometry. Poor separation of 93 kDa and 53 kDa peaks hinted at the presence of an intermediate peak. Indeed, deconvolution of the SEC profile revealed formation of a 2:1 (72 kDa) complex in addition to 2:2 (93 kDa) species, observed for the interaction of 14-3-3 with doubly phosphorylated STARD1 (Fig. 3D). The presence of the intermediary 2:1 peak could in addition explain the titration pattern observed in Fig. 3C. Overall, these results are in line with the optimization within the Ser57-containing motif, owing to the L59P substitution, indicating at the same time that Ser57 phosphosite contributes to the interaction with 14-3-3. The relatively efficient formation of 2:2 complexes in addition to 2:1 complexes indicates that the Ser195 phosphosite is not a significant contributor to the interaction. In any case, the presence of large amounts of unbound proteins during the SEC analysis, even upon the highest used concentrations, indicated weak affinity, requiring quantitative analysis.

**Fig. 3.**
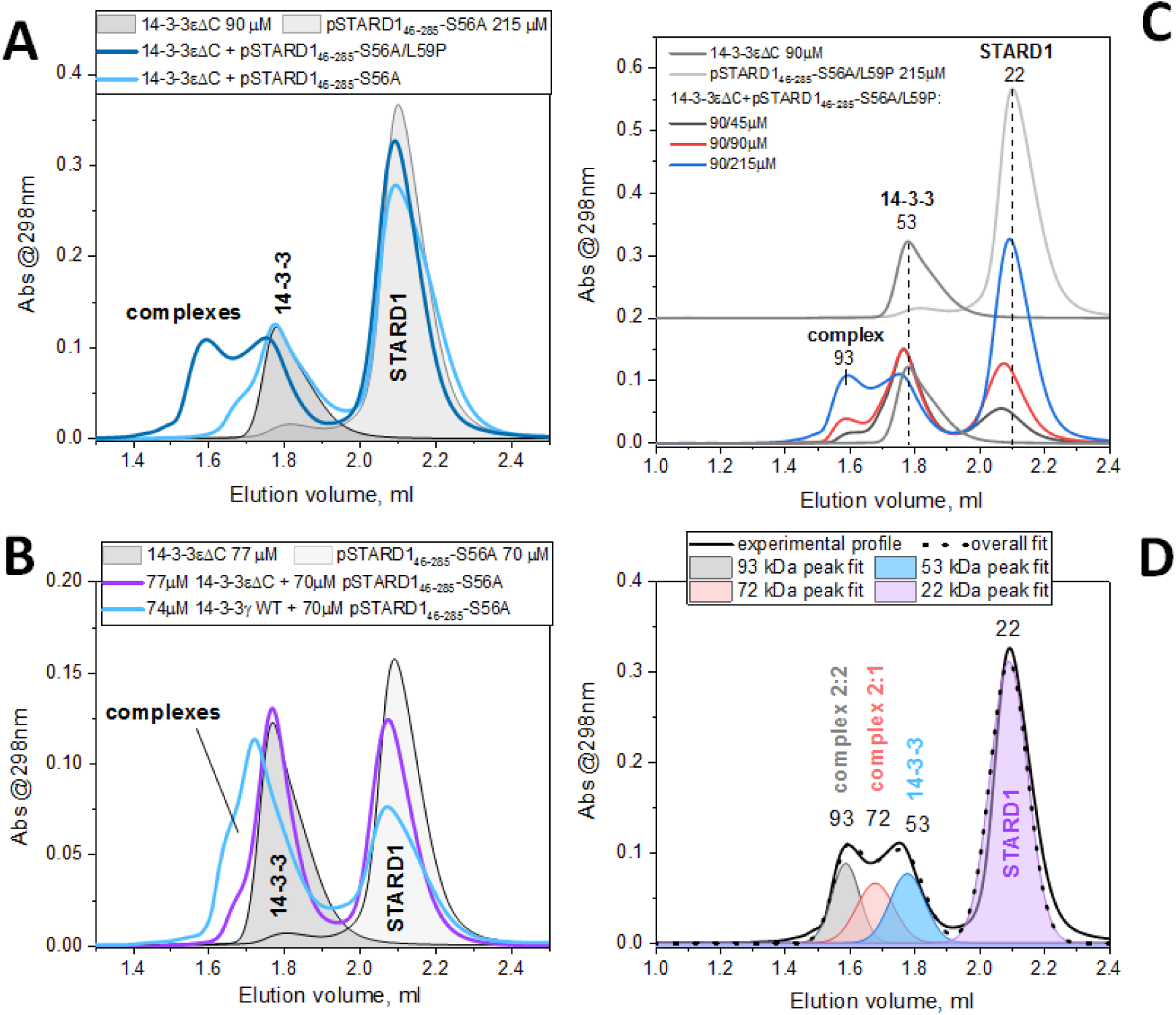
Interaction of STARD1 phosphorylated at both, Ser57 and Ser195, with 14-3-3. Interaction has been analyzed by size-exclusion chromatography using STARD1_46-285_-S56A and STARD1_46-285_-S56A/L59P constructs. The individual proteins or their mixtures at indicated molar concentrations were pre-incubated for over 20 min at room temperature before loading into Superdex 200 Increase 5/150 column at a 0.1 ml/min flow rate. Mw values, estimated from column calibration, are indicated in kDa. The peaks of the complexes are marked. Panel A shows comparison of the 14-3-3εΔC binding to phosphorylated STARD1_46-285_-S56A and STARD1_46-285_-S56A/L59P at a given concentration ratio (90 µM 14-3-3 and 215 µM STARD1, each per monomer). Panel B shows binding of the phosphorylated STARD1_46-285_-S56A to two different dimeric 14-3-3 constructs (molar concentrations per monomer are indicated). Panel C shows titration of 14-3-3εΔC by phosphorylated STARD1_46-285_-S56A/L59P (molar ratios are indicated). Panel D shows deconvolution of the chromatographic profile corresponding to a mixture of 90 µM 14-3-3εΔC and 215 µM of phosphorylated STARD1_46-285_-S56A/L59P allowing to detect two peaks of the complexes with different stoichiometries.

### Affinity between 14-3-3 and doubly phosphorylated STARD1

We used surface plasmon resonance (SPR) to quantify the interaction between doubly phosphorylated STARD1_46-285_-S56A/L59P and 14-3-3 immobilized on a chip using the N-terminal His-tag. Given the up-regulation of 14-3-3γ and 14-3-3ε isoforms (4 and 1.6 times, respectively) during the acute phase of steroidogenesis [41] and the unique preference of 14-3-3ε to heterodimerize with other 14-3-3 isoforms [61, 62], we produced the 14-3-3ε/γ heterodimeric species for further analysis (Fig. 3A). The obtained sensorgrams yielded an apparent K_D_ of 13 µM for 14-3-3ε and 15 µM for 14-3-3ε/γ (Fig. 3B and Table 1) (See Methods for details). The K_D_ values are in the same range as reported for the singly phosphorylated HSPB6 peptide RRApS^16^APL binding to 14-3-3 [63]. This indicates that, most likely, the only contributor to the interaction is the Ser57 site of pSTARD1 (RRApS^57^LPG). Analysis using STARD1_46-285_-S56A did not yield useful data due to limited sample solubility, however, given the SEC data (Fig. 3A, B), much larger K_D_ values could be expected in this case.

**Fig. 4.**
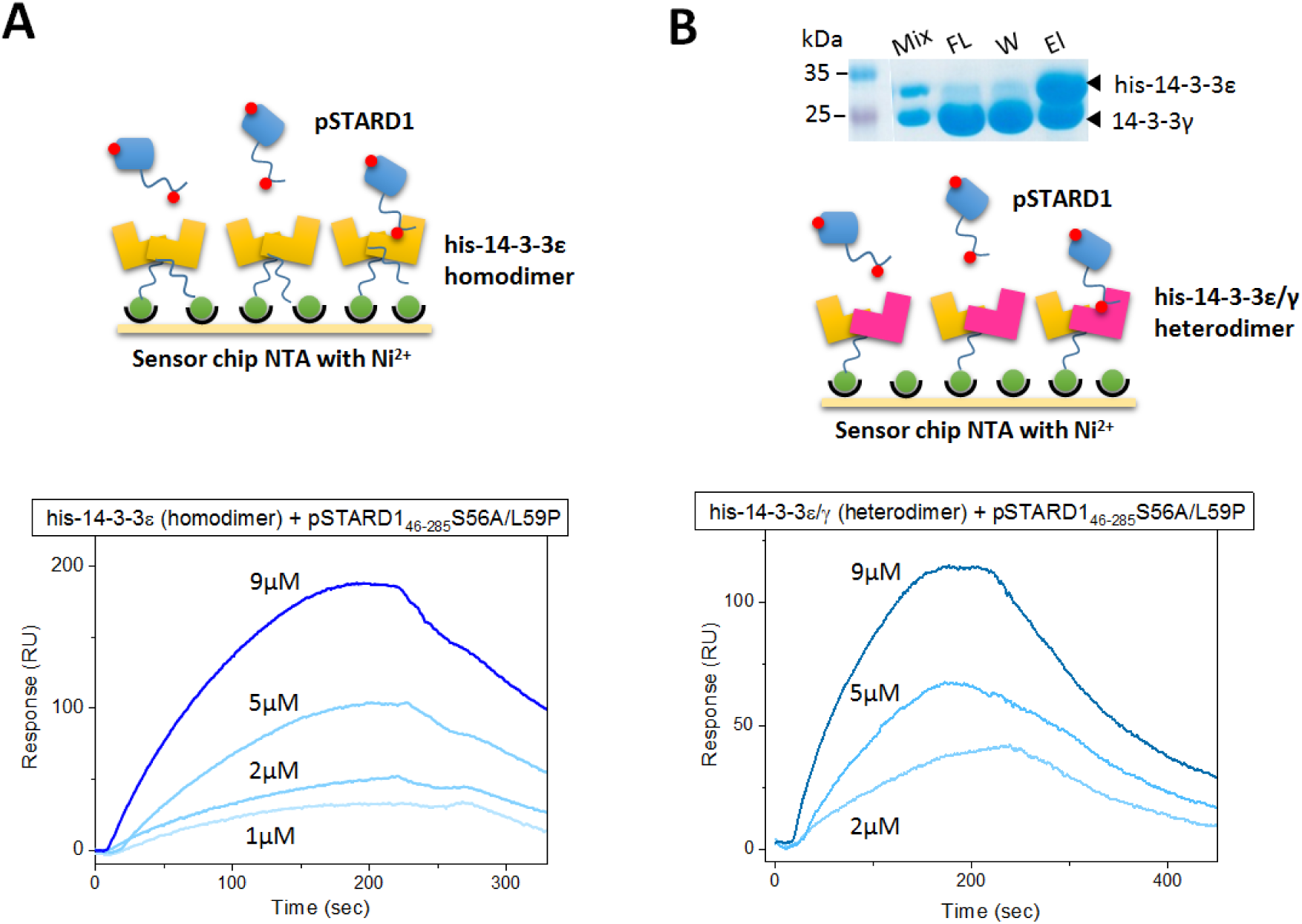
Interaction of doubly phosphorylated STARD1 with 14-3-3. Binding of STARD1_46-285_-S56A/L59P to either 14-3-3ε or its heterodimer with 14-3-3γ was analyzed by surface plasmon resonance. A. pSTARD1 binding to the His-tagged 14-3-3ε homodimer immobilized on a Ni-NTA chip. B. pSTARD1 binding to the heterodimer formed by the His-tagged 14-3-3ε and 14-3-3γ immobilized on a Ni-NTA chip. *Top*, schematic of the experiment; *bottom*, sensorgrams obtained for different flow concentrations of pSTARD1. Panel B also shows SDS-PAGE of the fractions obtained during preparation of the 14-3-3 heterodimer based on selective binding of His-tagged 14-3-3ε on the immobilized metal-affinity chromatography column and the unique preferential heterodimerization propensity of this isoform [62]. Mix – the initial mixture loaded on the IMAC column, Fl – flowthrough, W – washed fraction, El – eluate.

It is known that binding to 14-3-3 dimers is greatly enhanced for doubly compared to singly phosphorylated peptides [50]. In this respect, the affinity of the interaction measured by SPR (13-15 µM) was much lower than expected for binding using two phosphosites. This questioned whether both Ser57 and Ser195 sites contributed to interaction with the 14-3-3 dimer.

### Crystal structures of 14-3-3 complexes with STARD1 phosphopeptides

To obtain structural insights into the 14-3-3 interaction with STARD1 phosphopeptides, we exploited the recently introduced approach [60], which is particularly useful for transient and unstable interactions. To this end, we fused either the RRSS^57^LLGSR or RRGS^195^TCVLA peptide of STARD1 to the C terminus of the 14-3-3σΔC core via a GSGSL linker. This approach allows to enhance binding of phosphopeptides, while normally preserving its authentic binding conformation [60]. The resulting chimeric proteins containing the Ser57 and Ser195 peptides were co-expressed with PKA to ensure stoichiometric phosphorylation of the target sites, purified and their structures were determined by protein crystallography at 2.04 and 2.63 Å resolution, respectively (Supplementary Tables 2,3).

In the Ser57 peptide structure, only Ser57 is phosphorylated and bound in the amphipathic groove of 14-3-3. The traceable residues RRSpSLL adopt a conformation and a sophisticated network of polar contacts that are typical for the mode I peptides bound to 14-3-3 (Fig. 5), with the caveat that position +2 is occupied by the non-canonical leucine, which is expected to make the interaction less stable. Despite the relatively high resolution, three C-terminal residues of the peptide, GSR, are not seen due to disorder.

**Fig. 5.**
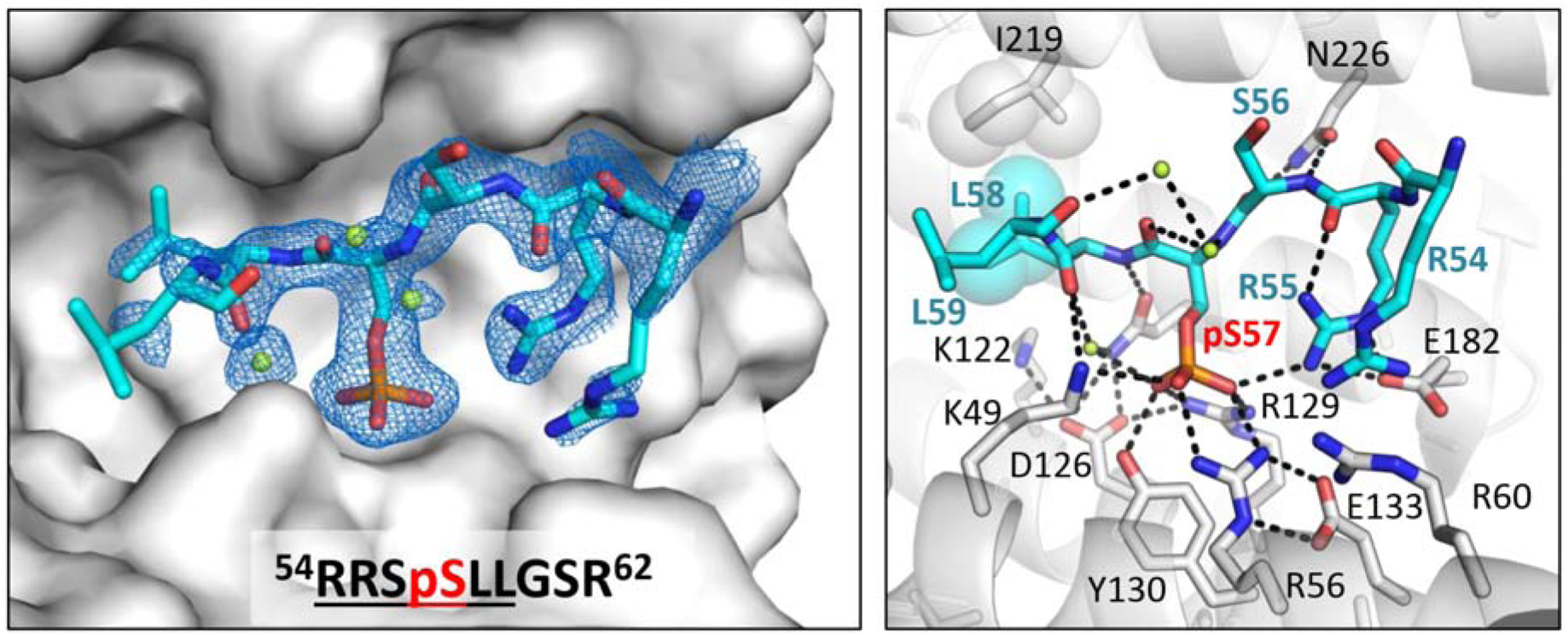
Molecular interactions of 14-3-3σ with the Ser57-containing STARD1 phosphopeptide revealed by 2 Å crystal structure. *Left*, 14-3-3 is shown as molecular surface (grey), peptide (atomic model with carbons in cyan, oxygens in red, nitrogens in blue and phosphate in orange) is shown with the corresponding 2F_o_-F_c_ electron density maps contoured at 1σ. Water molecules are in limon green. Residues of the phosphopeptide clearly visible in the electron density, are underlined in the sequence shown below. *Right*, A magnified view showing the network of 14-3-3:peptide interactions with several key 14-3-3 residues labelled in plain font and the peptide residues labelled in bold, with the central phosphoserine label in red. Polar contacts are shown by black dashed lines, hydrophobic contacts are shown by semitransparent spheres.

The structure for the protein with fused Ser195 peptide is unusual as the two 14-3-3 protomers bind the Ser195 peptides in two distinct conformations, an extended and a bent one, with all residues of the RRGpS^195^TCVLA peptide traceable (Fig. 6). In the first case, the adopted conformation displays the characteristic kink outward the amphipathic groove, despite Cys197 occupying the position of the canonical kink-promoting Pro residue at position +2. In the second case, the peptide adopts a turn in the polypeptide chain (Fig. 6). Although both conformations preserve contacts typical for the 14-3-3:phosphopeptide interactions, the extended conformation appears to be more advantageous due to additional hydrophobic contacts made by V198-L199 residues of the peptide (Fig. 6).

**Fig. 6.**
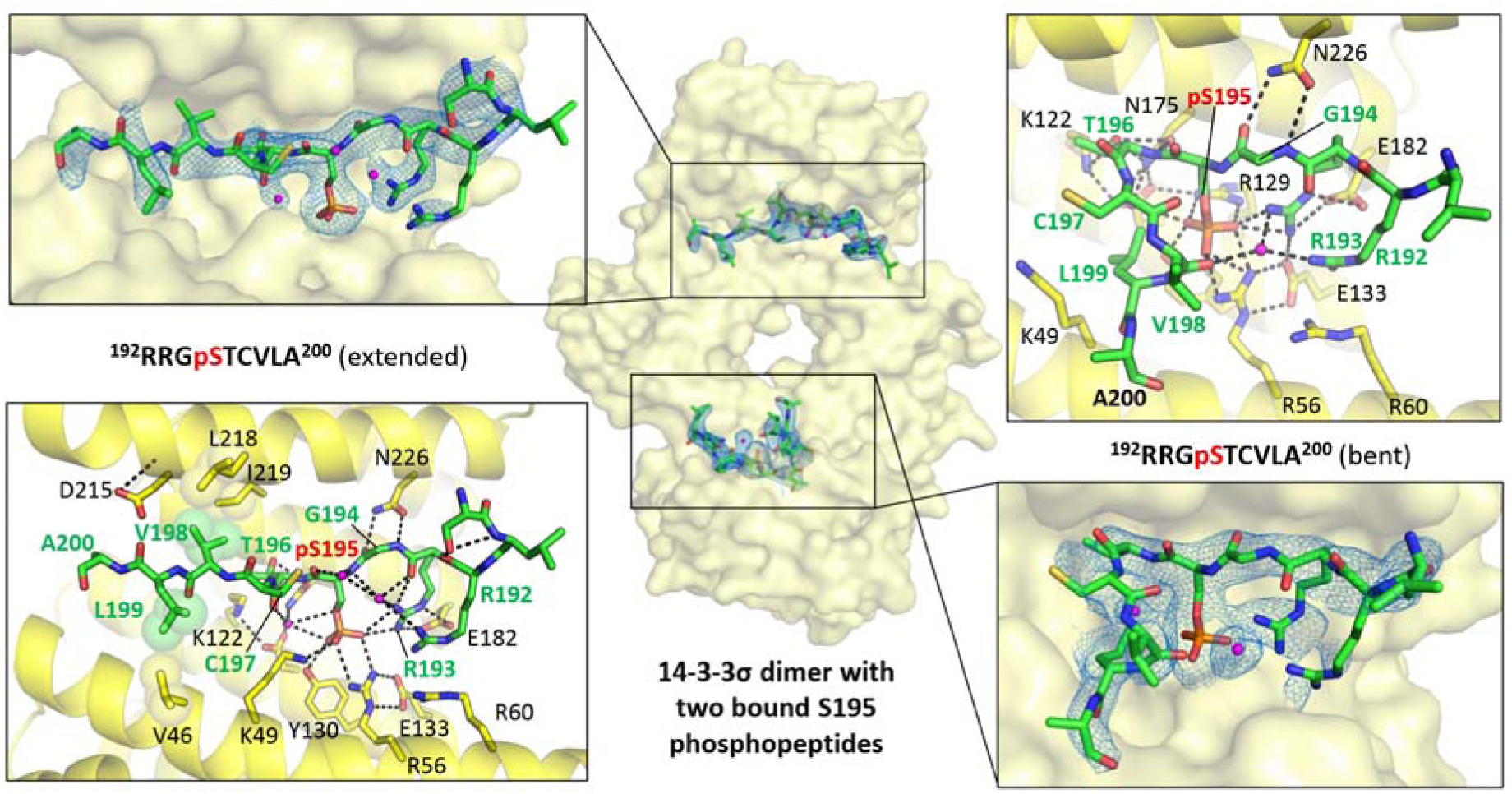
Molecular interactions of the 14-3-3 protein with the Ser195-containing STARD1 phosphopeptide. Crystal structure, derived at 2.6 Å resolution, revealed two distinct conformations of the phosphopeptide, with all residues traced for each conformation. *Middle*, the overall view of the 14-3-3 dimer with the two peptides occupying amphipathic grooves. *Left*, a closeup view showing molecular interactions for the extended conformation of the phosphopeptide. *Right*, a closeup view at the bent conformation of the phosphopeptide with carbon atoms in green, nitrogens in blue, oxygens in red, water molecules in magenta. 2F_o_-F_c_ electron density corresponding to the phosphopeptide is contoured at 1σ. Several key 14-3-3 residues are labelled in plain font and the peptide residues labelled in bold, with the central phosphoserine label in red. The phosphopeptide sequences are also depicted, with with the central phosphoserine label in red. Hydrophobic contacts involving the C-terminal V198-L199 residues of the phosphopeptide are shown by semitransparent spheres.

Notably, even the bent conformation of the Ser195-peptide observed in the present crystal structure (Fig. 6, right) is completely different to its β6-β7 loop conformation seen in the crystal structure of the stand-alone STARD1 domain (Fig. 1B and 7). In particular, Arg192 and Cys197 residues occupy the classic 14-3-3 binding determinants at positions -3 and +2 (Fig. 2A) [50]. It is clear that these residues cannot contribute to 14-3-3 binding in the context of the natively folded STARD1 conformation where they are part of a tight β6-β7 loop within the 9-stranded β-sheet (Fig. 1B).

**Fig. 7.**
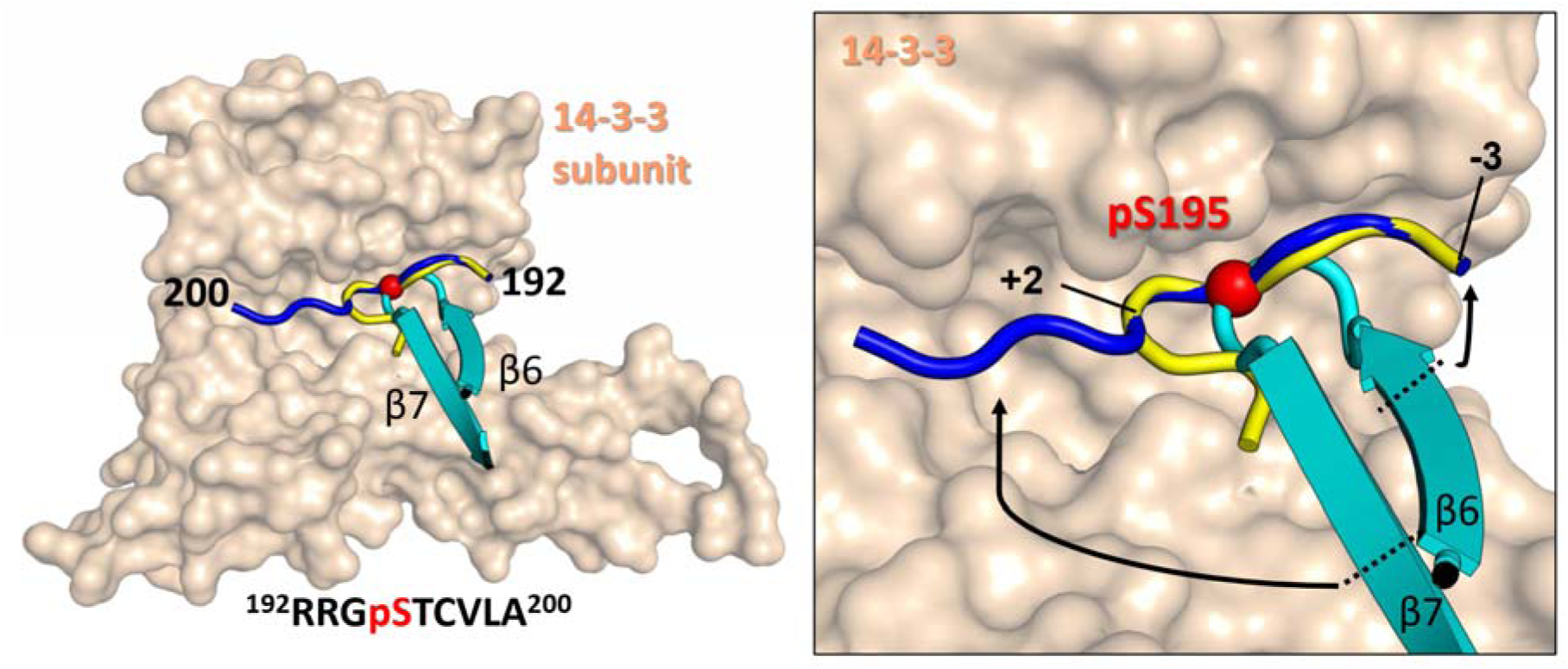
Comparison of the STARD1 Ser195 peptide conformation in the 14-3-3 bound and unbound states. Two conformations of the phosphopeptide observed in complex with 14-3-3 (Fig. 8) are overlaid with the conformation observed within the STARD1 structure where the phosphopeptide is part of the β6-β7 hairpin (PDB 3P0L), which suggests conditional binding to 14-3-3. *Left*, overall view at the 14-3-3 subunit with the three conformations of the Ser195-peptide shown in three different colors. The peptide sequence is shown below, phosphoserine-195 is in red. *Right*, a closeup view at the binding site showing tentative unfolding (black arrows) that is required for the β6-β7 loop to adopt an extended conformation. Cα atoms of the key 14-3-3 binding amino acids at positions -3 and +2 relative to the central phosphoserine (red) are indicated.

Of note, in structures of all 14-3-3 complexes with bound phosphopeptides or complete partner domains, phosphopeptides have relatively extended conformations, in line with the preferential binding of intrinsically disordered peptides [64]. Thus, in order to be bound in the amphipathic groove in a conformation typical for many phosphopeptide targets, the β6-β7 hairpin has to unfold and adopt the extended conformation (Fig. 7). These considerations are in line with our SEC data indicating that i) in their native conformation, neither STARD1 nor STARD3 domain show any appreciable binding to 14-3-3 (Supplementary Figs 1 and 2) and ii) that even the doubly phosphorylated STARD1 constructs display only moderate affinity towards 14-3-3 (Fig. 3 and 4). Interestingly, the 6-residue loop containing Ser195 appears flexible enough to allow for phosphorylation of this site within the folded recombinant STARD1 (Fig. 2D). The difference with respect to 14-3-3 is apparently explained by the fact that PKA recognizes RRXS substrate motif which is only part of the consensus sequences for 14-3-3 (Fig. 2A) and, therefore, has less strict structural restrictions. Indeed, it is known that even non-contiguous structural sites on the surface of globular proteins can serve substrates for efficient phosphorylation [65].

Since only the native conformation of the β6-β7 hairpin segment encompassing the Ser195 phosphosite would be compatible with the sterol binding, we decided to probe this by time-resolved anisotropy measurements using the fluorescently labelled cholesterol analog, 22NC.

### In complex with 14-3-3, doubly phosphorylated STARD1 retains native conformation capable of ligand binding

The kinetics of fluorescence anisotropy for free 22NC, 22NC in complex with phosphorylated STARD1_46-285_-S56A/L59P and 22NC in a larger complex between the 22NC-bound STARD1 and 14-3-3, were measured with picosecond time resolution (Fig. 8A). In common with other cholesterol derivatives carrying the NBD group [40], in the absence of proteins 22NC fluorescence was significantly quenched by solvent. Relaxation of 22NC fluorescence anisotropy was characterized by a 115 ± 23 ps time constant (Fig. 8B), which corresponds to quickly rotating solvatochromic fluorophores. Addition of 10-molar excess of phosphorylated STARD1_46-285_-S56A/L59P led to a 116-fold increase in the fluorescence signal and altered the kinetics of fluorescence anisotropy. The latter became clearly biphasic, with almost identical yield of fast (84 ± 6 ps, 48.0%) and slow (9.2 ± 0.2 ns, 52.0%) components (Fig. 8B). The appearance of the additional component with a longer correlation time of ∼9.2 ns can be explained only by formation of a 27 kDa complex of the 22NC with STARD1, which corresponds to a much slower rotation and has the predicted correlation time of 11 ns (See Methods for details). Although the presence of the fast component indicates that the 22NC fluorophore can rotate with a rate similar for the free dye, we assigned this component to 22NC accommodated within the cholesterol-binding cavity of STARD1, due to the following reasons. First, given the nanomolar ligand binding affinity [40] and the concentrations used (1 µM ligand and 10 µM STARD1), all 22NC molecules are expected to be bound to STARD1. Second, the solvatochromic property of 22NC would mean ∼116 times lower fluorescence signal from the dissociated dye that could not be detected in the presence of the dominating protein-bound form. In the presence of an excess of 14-3-3γ, we observed the biphasic kinetics of fluorescence anisotropy, with the fast (72 ± 6 ps, 51.4%) and slow (40.0 ± 0.8 ns, 48.6%) components (Fig. 8B). Importantly, the long correlation time significantly increased compared to that in the absence of 14-3-3 (9.2 ns), which can be explained by formation of a larger protein complex embedding 22NC that further restricts its rotation speed. The experimental value of 40 ns agrees well with the correlation time of 35 ns predicted for the 85 kDa 14-3-3γ/pSTARD1/22NC 2:1:1 complex that is expected to be a dominant 22NC-containing species at concentrations used. Difference between the observed (40 ns) and predicted (35 ns) species can be explained by a non-spherical shape of the complex (Fig. 8A). As in the STARD1/22NC complex, the presence of the fast component (72 ± 6 ps) indicates that 14-3-3 binding does not affect the ability of the fluorophore to rotate within the cholesterol-binding cavity. The formation of the larger 22NC complex with STARD1 and 14-3-3 appears to preserve the rotational freedom of the solvatochromic NBD group in the bound ligand, which explains fast depolarization of NBD fluorescence.

**Fig. 8.**
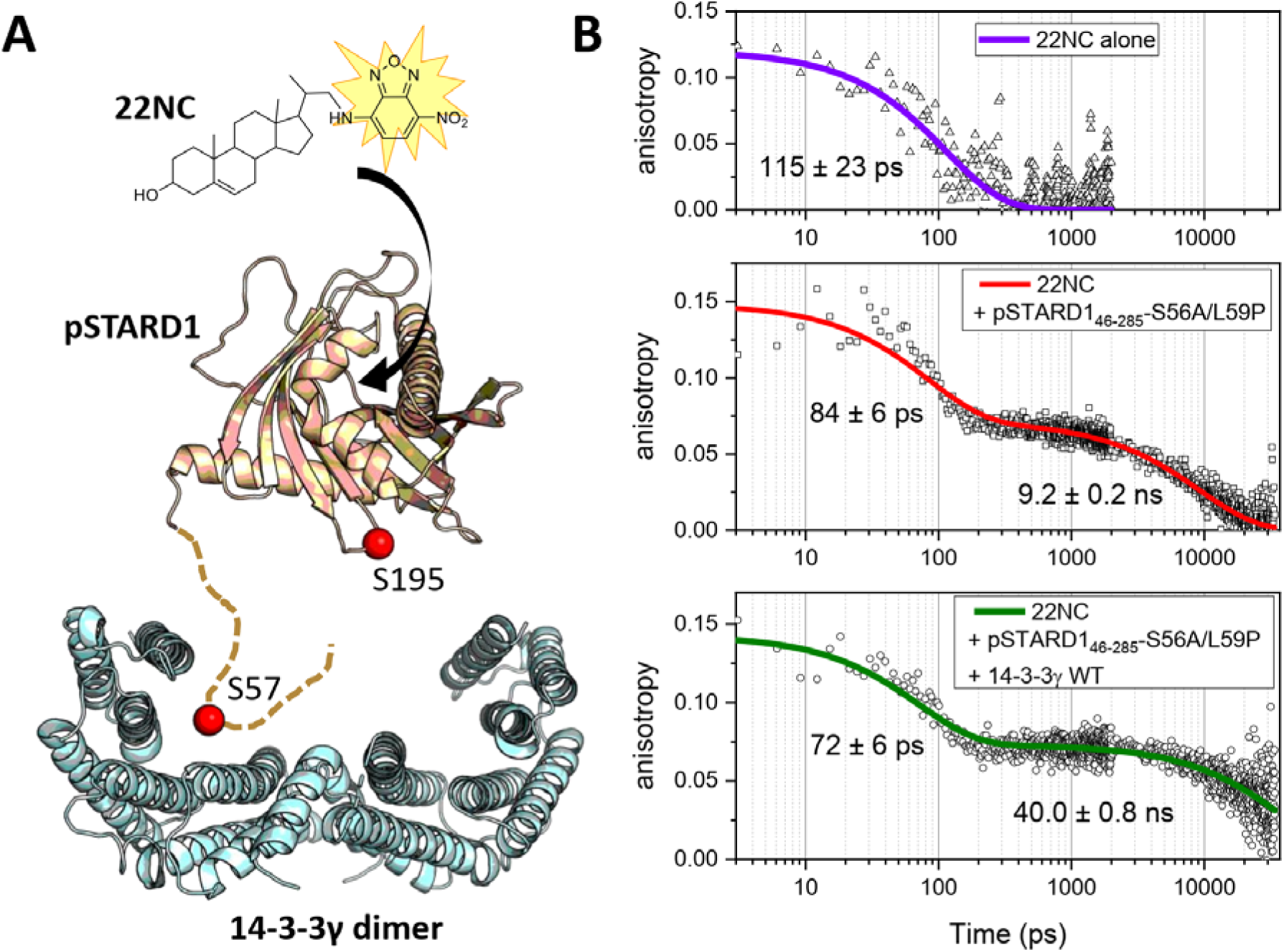
STARD1_46-285_-S56A/L59P phosphorylated at Ser57 and Ser195 retains ligand-binding ability upon interaction with 14-3-3_γ_. A. Graphical abstract of the experiment. NBD-labeled cholesterol analog 22NC (1 µM) was supplemented with 10:1 molar excess of phosphorylated STARD1_46-285_-S56A/L59P (10 µM) and then wild type 14-3-3γ (100 µM) was added at further 10-fold excess, to ensure complete binding of the 22NC reporter. Protein partners are shown in scale using PDB 3P0L (STARD1) and 6S9K (14-3-3γ). B. The kinetics of the 22NC fluorescence anisotropy was recorded with single-photon counting and picosecond temporal resolution for free 22NC (top graph), its complex with pSTARD1 (middle) and within the 14-3-3γ-bound pSTARD1 (bottom). Numbers indicate corresponding correlation times.

Together these data demonstrate that in complex with 14-3-3, the phosphorylated STARD1 retains its ligand-binding ability and conformation. This observation disproves the notion that the Ser195-phosphosite contributes to the interaction with 14-3-3. Such interaction would require significant structural rearrangements in STARD1 which could, in principle, take place *in vivo*, for example during the proposed molten-globule-like structural transformation of STARD1 or when it undergoes unfolding accompanying its import into mitochondria.

## DISCUSSION

In spite of critical importance for physiology and therapy, the mechanism of regulation of steroidogenesis by STARD1 remains puzzling. Upon tropic hormone stimulation, this protein is expressed *de novo*, targeted to the mitochondria and rapidly accumulated as phosphoprotein [3, 4, 11, 35]. STARD1 appears to cooperate with other proteins at the outer mitochondrial membrane (OMM) [66] (Fig. 9, **step 1**), where its phosphorylation in response to elevated cAMP occurs with the help of anchored regulatory subunits of PKA [38, 39, 67]. Recent data indicate that members of the 14-3-3 protein family, up-regulated in the acute phase of steroidogenesis [41], can partner with STARD1 (Fig. 9, **step 2**). It was proposed that, depending on phosphorylation of STARD1 and 14-3-3 and on the oligomeric state of 14-3-3, 14-3-3 can negatively affect steroidogenesis by binding to STARD1 [41, 42]. Intriguingly, STARD1 contains at least two putative 14-3-3 binding motifs centered at Ser57 and Ser195, that are readily phosphorylated by PKA ([32] and this work). In addition, it has been reported that STARD3, a close paralog of STARD1 possessing the same fold of its cholesterol-binding domain and responsible for placental steroidogenesis [11, 22], can interact with 14-3-3 proteins in a phosphorylation-*independent* manner using its KSASNP motif [48].

**Fig. 9.**
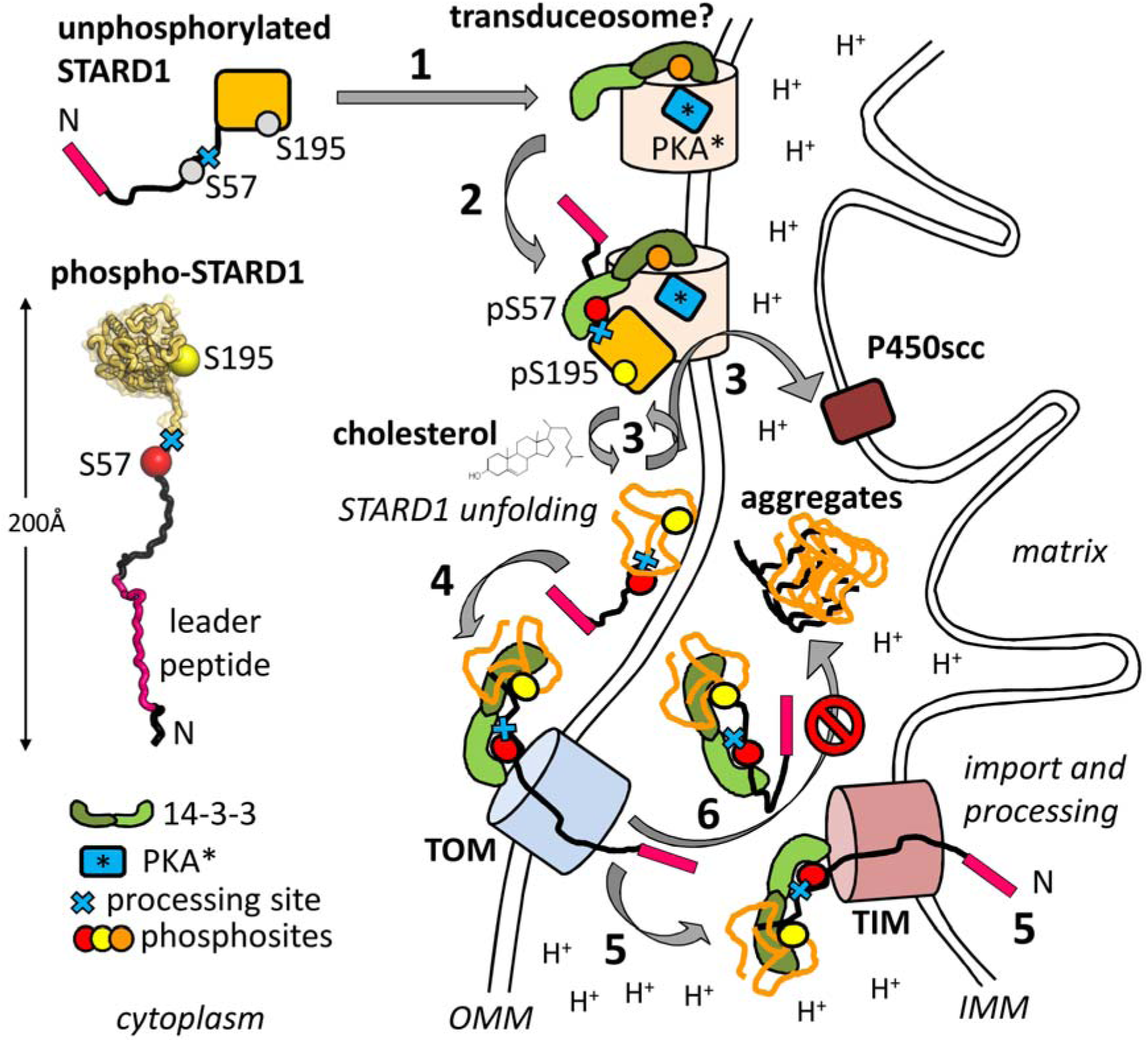
Plausible mechanism of STARD1 regulation by the 14-3-3 protein. Key elements of each protein are labelled. Insert shows the full STARD1 model (residues 1-285) built by Bioserf [75] and depicted schematically to show the real proportion of the N-terminal peptide and STARD1 domain. Steps 1-6 are described in the text. IMM – inner mitochondrial membrane, OMM – outer mitochondrial membrane, TIM and TOM – translocase components of the inner and outer membrane, respectively. Putative protein complex known as transduceosome [66] is marked by a question mark and symbolizes OMM proteins recruiting STARD1 and 14-3-3 (including AKAP121 and PKA for STARD1 phosphorylation on site [39]). Protons illustrate low pH in the intermembrane space, which may provoke STARD1 unfolding and aggregation, counteracted by 14-3-3. Ser195 embedded within the β-sheet of native STARD1 is not recognized by 14-3-3 but likely becomes operational upon STARD1 unfolding in vicinity to the mitochondrial membrane or within the intermembrane space [19, 24] (steps 4 and 6). Bidentate binding of unfolded STARD1 phosphorylated at both, Ser57 and Ser195 sites, to the 14-3-3 dimer protects STARD1 from dephosphorylation, aggregation and degradation, interfering with STARD1 import into mitochondria and its processing. Conditional binding of Ser195 phosphosite to 14-3-3 can influence the folded-unfolded STARD1 turnover required for the cholesterol transfer.

Despite intensive research, many questions remained about the molecular mechanism of STARD1 regulation, prompting us to dissect the role of the START domain and the two principal phosphorylation sites present in STARD1, in interaction with the 14-3-3 protein. This analysis was carried out using soluble versions of STARD1 lacking part of the N-terminal peptide responsible for mitochondrial localization.

Our data show that despite the presence of the previously reported 14-3-3 binding elements such as the phosphosite around Ser195 in STARD1 and the KSASNP motif in STARD3, isolated START domains of STARD1 and STARD3 do not form stable complexes neither with dimeric nor monomeric forms of 14-3-3, even at the high protein concentrations (Supplementary Fig. 1 and 2). However, stepwise extension of the STARD1 construct along the N terminus (Fig. 2) led to detectable interaction. The extended STARD1 construct that interacted with 14-3-3 was phosphorylated at both Ser57 and Ser195 positions, with the interaction observed only for the dimeric 14-3-3 proteins (Fig. 3), and not the monomeric 14-3-3 mutant m58E protein (data not shown). While optimization of the Ser57-peptide by L59P amino acid substitution enhanced interaction with 14-3-3, the Ser195 phosphosite residing in the native START domain did not appear to contribute into 14-3-3 binding. As a result, the micromolar binding affinity of the doubly phosphorylated native STARD1 to 14-3-3 (13-15 µM) was similar to that of a singly phosphorylated peptide with a similar sequence (Fig. 3-4). Crystallographic analysis of molecular interactions with 14-3-3 revealed canonical conformations for both Ser57 and Ser195 phosphopeptides (Figs 5-6).

The Ser57 phosphopeptide (<u>RRSpS^57^LL</u>GSR) of STARD1 is expected to be intrinsically disordered (Fig. 1) and appears to be a critical 14-3-3-binding site. Notably, almost identical phosphopeptides, <u>RRSpS^87^LL</u>SRS in the human proapoptotic Bcl-2-like protein 11 (Uniprot O43521) and <u>RRSpS^435^LL</u>SLM in the RAB11 family interacting protein 1 (Uniprot Q6WKZ4), can likewise be recognized by 14-3-3. Based on identical core residues underlined in the sequences above, it can be expected that primary interactions of these two proteins with 14-3-3 are the same as seen in the crystal structure of the phospho-Ser57 peptide of STARD1, reported here. Together, the three 14-3-3 interaction partners represent a rare example of proteins with almost identical 14-3-3-binding segments.

The biological role of the Ser195 phosphopeptide (RRGpS^195^TCVLA) is intriguing. Although it has a suboptimal 14-3-3 binding sequence, it is well-defined in crystal structures. It is able to adopt at least three different conformations: the two distinct, extended conformations reported here and as part of a rigid β6-β7 hairpin in the intact STARD1 (PDB 3P0L) [13]. According to the gatekeeper hypothesis introduced by Yaffe [68], a doubly phosphorylated polypeptide containing even suboptimal 14-3-3 binding sites could lead to a highly cooperative conditional binding owing to an increased local concentration of the second binding site (Ser195) following the primary Ser57-site binding. Nevertheless, we could not detect the expected, high-affinity interaction with 14-3-3 for the phosphorylated STARD1 containing START domain capable of binding fluorescently labeled cholesterol analog 22NC (Fig. 8). The extended conformations of the Ser195 peptide observed in the crystal structure (Fig. 6) strongly indicate that the Ser195 phosphosite can contribute to 14-3-3 binding only upon partial unfolding of the β-sheet that comprises the Ser195 phosphopeptide. Hence, the folded state of the START domain prevents it from adopting alternative conformation necessary for binding to 14-3-3 (Fig. 7). To our knowledge, such conditional binding to 14-3-3 proteins has not been structurally characterized before. This precedent warns prediction of 14-3-3 binding sites based on simple comformity to 14-3-3 binding consensus motifs and location in the intrinsically disordered regions, as cryptic 14-3-3 binding sites can most likely exist in other proteins.

Based on the available evidence, we hypothesize that the Ser57 phosphosite plays a role in the 14-3-3-mediated attachment of STARD1 to transduceosome (Fig. 9, **step 2**). The residence of STARD1 at the OMM likely involves molten-globule structural transition [19, 24]. In so far poorly understood cooperation with other proteins at the OMM, this enables repeated cholesterol binding and release (Fig. 9, **step 3**), to ensure cholesterol delivery to the cholesterol side-chain cleavage cytochrome P450 (P450scc) (Fig. 9, **step 3**), which converts it into pregnenolone [1]. Since Ser57 is proximal to the predicted sites of STARD1 processing (residues 55-56 [30]), 14-3-3 binding to this site in intact STARD1 would also regulate its mitochondrial import (Fig. 9, **steps 4 and 5**). Indeed, 14-3-3 binding is known to inhibit proteolytic degradation, dephosphorylation and aggregation of target proteins [69-71] (Fig. 9, **step 6**). Therefore, 14-3-3 interaction with phosphorylated Ser57 would sequester STARD1 in the cytoplasm by interfering with its mitochondrial import and processing. This, in turn, can prolong the action of STARD1 as a key factor in cholesterol transport, during the acute phase of steroidogenesis, in line with the ‘pause-transfer’ hypothesis. It is pertinent to note that 14-3-3 also interacts with the phosphorylated mitochondrial import sequence of the iron-sulfur cluster assembly enzyme [72]. Our results support the notion that the Ser195 phosphosite can also contribute to binding to the 14-3-3, but only upon STARD1 unfolding [41] (Fig. 9, **steps 4-6**). This mode of binding is relevant to the reported chaperone-like, anti-aggregation activity of 14-3-3 [71], and can accompany structural transition of STARD1 into a molten-globule conformation [19, 21, 24] (Fig. 9, **step 3**). Such interaction may also occur upon STARD1 unfolding during the mitochondrial import (Fig. 9, **steps 4 and 5**). In accordance, 14-3-3 proteins were proposed to play a role of mitochondrial import stimulating factors as part of the so-called guidance complex including chaperones such as HSP70 [73]. In turn, such unique, conditional binding using Ser195 phosphosite would also affect the rate of transition of STARD1 between folded and unfolded states and may interfere with STARD1 degradation and dephosphorylation (Fig. 9, **steps 3-6**).

The reported 50% enhancement of the steroidogenic activity of STARD1 by Ser195 phosphorylation [32] and the inability of S195A mutated STARD1 transgene to rescue STARD1^-/-^ phenotype in knockout mice [9] together hint at the critical importance of this phosphorylation site in STARD1 functioning. In addition, S195A mutation was found in a patient with classic LCAH [9]. Individuals with the nonclassic LCAH disorder may have the R192C mutation associated with ∼2-fold reduction in the steroidogenic activity [74]. Since this mutation is at the key -3 position within the Ser195 phosphopeptide of STARD1, it is likely it will affect interaction with 14-3-3 and hence interfere with STARD1 function and mitochondrial import, a hypothesis that warrants further investigation *in vivo*.

## METHODS

### Cloning, protein expression and purification

Wild-type full-length human 14-3-3γ (Uniprot P61981) and human 14-3-3ε devoid of the C-terminal flexible tail (14-3-3εΔC construct; Uniprot P62258) were produced as described [63, 76]. The engineered monomeric form of the human 14-3-3ζ (Uniprot P63104) carrying the monomerizing amino acid substitutions ^12^LAE^14^ → ^12^QQR^14^ on top of the dimer-destabilizing phosphomimicking mutation, S58E, was produced as described [57].

Chimeric constructs of the human 14-3-3σΔC with the STARD1 phosphopeptides Ser57 and Ser195 were produced on the basis of the previously described 14-3-3σ chimeric protein with the HSPB6 phosphopeptide [60], using high-fidelity *Pfu* polymerase and the following reverse primers: 5’-ATAT<u>CTCGAG</u>TCAACGAGATCCCAGCAGGCTGCTGCGGCGCAGGGATC-3’ for Ser57 and 5’-ATAT<u>CTCGAG</u>TCAAGCCAGCACACAGGTGGAGCCGCGGCGCAGGGATC-3’ for Ser195 peptides (*XhoI* sites are underlined). PCR products were cloned into a pET28-his-3C vector using *NdeI* and *XhoI* sites to produce proteins with the 3C protease-cleavable N-terminal His-tags [59, 60]. Such chimeric protein constructs contained amino acid substitutions ^75^EEK^77^ → ^75^AAA^77^ promoting crystallization [60]. Chimeric proteins were co-expressed with the catalytically active subunit of mouse PKA [59, 60] and purified by subtractive immobilized metal-affinity chromatography including 3C cleavage of the His-MBP tag and gel-filtration.

The 14-3-3γ/ε heterodimer was obtained as described [77]. Briefly, the His_6_-tagged 14-3-3ε was allowed to equilibrate with the excess of untagged 14-3-3γ (overnight at 4 °C) and then loaded on a 1 ml HisTrap HP column (GE Healthcare) while the flowthrough fraction containing 14-3-3γ was collected. After a washing step, the bound heterodimer was eluted with 200 mM imidazole and then buffer-exchanged into SEC buffer.

Human STARD1_66-285_ (Uniprot P49675) was cloned into the H-MBP-3C vector using *BamHI* and *HindIII* sites, expressed and purified as described [26]. The S195E phosphomimicking mutant of STARD1_66-285_ was obtained using the wild-type construct as a template, as described earlier [40, 54]. The longer constructs extending to residue 53 or 46 were obtained using the wild-type STARD1_66-285_ plasmid as a template, high-fidelity *Pfu* polymerase and the following forward primers: 5’-TATA<u>GGATCC</u>AGGAGGAGGTCCTCCCTGCTGGGAAGCAGGCTCGAGGAAACTCTCT ACAGTGACCAGG-3’ for STARD1_53-285_; 5’-ATATA<u>GGATCC</u>ACTTGGATTAATCAAGTTAGGAGGAGGGCTTCCCTGCTGGGAAGC-3’ for STARD1_46-285_-S56A and 5’-ATATA<u>GGATCC</u>ACTTGGATTAATCAAGTTAGGAGGAGGGCTTCCCTGCCGGGAAGC AG-3’ for STARD1_46-285_-S56A/L59P (*BamHI* sites are underlined). Codon-optimized full-length STARD1 sequence with the S56G mutation mimicking the sequence of bovine (or sheep) ortholog and flanked by *BamHI* and *HindIII* sites for subsequent cloning into the H-MBP-3C vector was ordered as gblocks from ITDNA Technologies (https://eu.idtdna.com/pages).

Human STARD3_216-444_ (Uniprot Q14849) was kindly provided by Prof. T. Friedrich (TU Berlin) and was cloned into the pQE80 vector to produce the His-tagged protein with the N-terminal sequence MRGSHHHHHHGSACEL**G**^**216**^**SDN..LGAR**^**444**^ (STARD3 sequence is in bold font). All constructs were verified by DNA sequencing in Evrogen (https://evrogen.com) and used to transform chemically competent cells of *Escherichia coli* BL21(DE3) already carrying pACYC-PKA plasmid for protein phosphorylation [59]. Purification of STARD1 constructs was achieved by subtractive immobilized metal-affinity chromatography including 3C cleavage of the His-MBP tag, and gel-filtration. The STARD3 construct retained the N-terminal his-tag. Phosphorylation of specific sites was analyzed by native gel-electrophoresis, gel band excision, in-gel trypsinolysis and tandem mass-spectrometry using a MALDI TOF/TOF ultrafleXtreme instrument (Bruker, Germany). To verify stoichiometric phosphorylation, phosphorylated STARD1 samples were incubated with either PKA or phosphatase (FastAP) in vitro followed by native PAGE analysis as described [59, 60]. Protein concentration was determined spectrophotometrically at 280 nm on a N80 Nanophotometer (Implen, Germany) using sequence-specific extinction coefficients calculated using the ProtParam tool in ExPASy.

### Analytical size-exclusion chromatography

The oligomeric state of proteins and interaction between 14-3-3 and STARD1 (or STARD3) constructs were analyzed by size-exclusion chromatography on a Superdex 200 Increase 5/150 column (GE Healthcare) operated at 0.1 ml/min flow rate using a Varian ProStar 335/ProStar 363 system. Where specified, a Superdex 200 Increase 10/300 column (GE Healthcare) operated at a 1.5 ml/min flow rate was used. The columns were equilibrated by a SEC buffer (20 mM Tris-HCl buffer, pH 7.6, containing 150 mM NaCl, 5 mM MgCl_2_ and 3 mM β-mercaptoethanol (ME)) and calibrated by the following protein markers: BSA trimer (198 kDa), BSA dimer (132 kDa), BSA monomer (66 kDa), ovalbumin (43 kDa), α-lactalbumin (15 kDa). The profiles were followed by 298 nm absorbance to avoid detector overload by highly concentrated protein samples. All experiments were performed at least three times and the most typical results are presented.

### Small-angle X-ray scattering (SAXS)

The solution structure of STARD3 was analyzed by SAXS at the P12 beamline (PETRA III, DESY Hamburg, Germany) using inline SEC for the peak-specific data collection, to avoid contamination with protein oligomers and aggregates. The 100-µl sample (8 mg/ml) was loaded on a Superdex 200 Increase 10/300 column (GE Healthcare) pre-equilibrated with filtered and degassed SEC buffer containing 3% glycerol and 5 mM dithiothreitol instead of MgCl_2_ and β-mercaptoethanol. The column was operated at a 0.5 ml/min flow rate. Only the monomeric STARD3 peak was used for analysis. SAXS data frames were buffer subtracted and processed using CHROMIXS [78]. No radiation damage for protein frames was observed. The final SAXS curve averaged across the STARD3 peak was used for modeling in CORAL [79]. For this, the 5I9J crystal structure of STARD3 was supplemented with 31 flexible N-terminal residues to account for differences in the constructs. CRYSOL [80] was exploited to calculate the theoretical scattering profile from models with varying conformation of the N-terminal peptide as well as the discrepancies between the profiles calculated for each conformation and the experimental SAXS profile.

### Crystallization and structure determination of the chimeric 14-3-3/STARD1 phosphopeptide protein constructs

Search of suitable crystallization conditions for the chimeric 14-3-3 proteins containing the STARD1 phosphopeptides was performed immediately after the final purification step, by the sitting-drop vapor diffusion, using commercial screens: JCSG+ (Molecular Dimensions), Index, Crystal Screen (Hampton Research) and PACT (Qiagen). Crystals formed in several conditions with the best conditions summarized in Supplementary Table 2. Crystallization plates were stored at 6 ′C and were periodically monitored using a Rigaku plate imager equipped with a Vis/UV-scanning and detection system.

Crystals were mounted in nylon loops directly from the crystallization drops and vitrified in liquid nitrogen. X-ray diffraction data (Supplementary Table 3) were collected at 100 K at I04 and I04-1 beamlines at the Diamond Light Source (UK) using Dectris EIGER2 × 16M (I04) and PILATUS 6MF (I04-1) detectors.

Diffraction data were processed using XDS/XSCALE [81]. The structures were solved by molecular replacement using MolRep [82] with 14-3-3σ dimer (PDB 5LU1) as a search model. The phosphopeptides were built *de novo* by manual placement of the corresponding amino acid residues into difference electron density maps in Coot [83]. Structure refinement was conducted using Buster 2.10.3 [84] which included a rigid-body refinement of all chains followed by an all-atom and individual B-factor restrained refinement (the refinement statistics is shown in Supplementary Table 3).

### Surface plasmon resonance

The kinetics of pSTARD1_46-285_-S56A/L59P interaction with either the His-tagged 14-3-3ε or its heterodimer with untagged 14-3-3γ was studied by surface plasmon resonance on a BIAcoreTM X instrument (Pharmacia Biosensor AB, Sweden) using an NTA sensor chip (BR-1000-34) with a nitrilotriacetic acid (NTA) modified surface. One of the two cells was used as an analytical cell, with the second cell used as a reference. His_6_-tagged 14-3-3 was immobilized on the chip by passing its 1 mg/ml solution in a 20 mM Tris-HCl buffer, pH 7.5, 250 mM NaCl, 2 mM β-mercaptoethanol, 5 mM MgCl_2_ over the chip surface (total volume of 80 μl at a flow rate of 10 μl/min). The solution was passed only through the analytical cell, the reference cell remained closed. Then, both cells were opened and pSTARD1 samples with varying protein concentration were applied while recording the kinetics of protein interaction. Data processing and analysis were performed using BIAevaluation (Biacore) and OriginLab 9.0. Binding constants were calculated in the approximation of equilibrium conditions using equation:

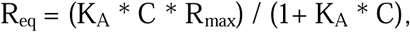

where K_A_ is the equilibrium binding constant, C is the concentration of the added protein, R_eq_ is the signal difference in the analytical cell and the reference cell at a given protein concentration, R_max_ is the limiting level of binding.

In the Scatchard coordinates (1/R_eq_ versus 1/C), this dependence takes a linear form. The intersection point of the straight line with the ordinate axis gives 1/R_max_, and the slope of the linearized dependence gives 1/K_A_*C*R_max_, whence the K_A_ value was obtained. The kinetic dissociation constant was calculated in the approximation of the irreversible reaction, because dissociation products are washed out of the reaction medium by a liquid flow and the reaction becomes irreversible.

The model of complex dissociation was calculated in BIAevaluation according to the equation:

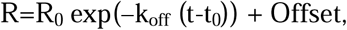

where R is the signal value at time t, k_off_ is the kinetic dissociation constant, Offset is the signal level at the initial time t_0_.

R and R_0_ values obtained from the dissociation region of the sensorgrams led to the k_off_ value determination. The relations K_A_ = 1/K_D_ = k_on_/k_off_ were used to determine the values of the kinetic association constant (k_on_) and the equilibrium dissociation constant (K_D_).

### Time-resolved fluorescence anisotropy

22NC cholesterol analog was dissolved in 96% ethanol to get a 200-µM stock solution (concentration was measured by absorbance at 470 nm using the extinction coefficient of 21,000 M^-1^ cm^-1^) and then diluted down to 1 µM into 20 mM Tris-HCl buffer, pH 7.5, containing 150 mM NaCl and 5 mM MgCl_2_.

22NC fluorescence in different environments (free dye or its complex with pSTARD1_46-285_-S56A/L59P in the absence or presence of 14-3-3γ) was recorded using a time- and wavelength-correlated single-photon counting (TCSPC) system based on a single TCSPC module (SPC-130EM, Becker & Hickl GmbH, Germany). Fluorescence was detected by a cooled ultrafast single-photon counting detector (HPM-100-07C, Becker and Hickl, Germany) with a low dark count rate (∼10 counts per second), coupled to a monochromator (ML-44, Solar, Belarus). Excitation was performed with a 455 nm laser diode (InTop, Russia) driven at a repetition rate up to 50 MHz. A 500-nm long-pass filter (Thorlabs, USA) was used to block excitation light. To measure the kinetics of fluorescence anisotropy we used a filter-based system with a set of two ultra-broadband wire-grid polarizers WP25M-UB (Thorlabs, USA) mounted into motorized rotation mounts K10CR1/M (Thorlabs, USA). Fluorescence decay kinetics *I(t)* were measured at different positions of the emission polarizer – in parallel (‖) and perpendicular (⊥) orientation to the excitation polarizer and the anisotropy kinetics *r*(*t*) was calculated as:

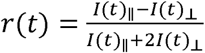

By using two different repetition rates of the laser (50 and 25 MHz) we achieved the time resolution of up to 3.3 ps and were able to follow slow changes in the anisotropy of the dye in complex with proteins. The temperature of the sample was stabilized at 25 °C by a Peltier-controlled cuvette holder Qpod 2e (Quantum Northwest, USA). Analysis of fluorescence and anisotropy decay curves was performed by SPCImage (Becker and Hickl, Germany) and Origin Pro 9 (OriginLab Corporation, USA) software packages. The experiments were performed at least three times.

The correlation time 0 (for the rotation of a spherical protein) was calculated according to Perrin equation [85]:

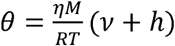

where *η* is the viscosity of the solvent, *M* is the molecular weight of the protein, *v* and *h* are the partial specific volume of the protein and its degree of hydration (both in ml/g), respectively. For estimations, water viscosity *η* was taken as 0.9 cP (at 25 °C), partial specific volume and the degree of hydration were taken as 0.73 ml/g and 0.4 ml/g [86], respectively.

## Data availability

Atomic coordinates and structure factors were deposited with the PDB under accession codes 6T5H and 6T5F. All other data supporting the findings of this study are available from the corresponding author upon reasonable request.

## Acknowledgments

N.N.S. is thankful to Prof. T. Friedrich (TU Berlin) for providing the STARD3 plasmid, to Dr. Ya. Faletrov (Minsk, Belarus) for providing 22NC and to Dr. Cy Jeffries (EMBL Hamburg) for help during the SAXS-756 experiment. We would like to thank Sam Hart and Johan Turkenburg for help during data collection, the Diamond Light Source for access to synchrotron beamlines (Proposal No. mx-13587). Mass-spectrometry analysis was carried out at the Shared-Access Equipment Centre “Industrial Biotechnology” of the Federal Research Center “Fundamentals of Biotechnology” of the Russian Academy of Sciences. This work was supported by the Russian Science Foundation (grant 19-74-10031 to N.N.S.) and the Wellcome Trust (206377 award to A.A.A.). Protein-protein interactions were studied in the framework of the Program of the Russian Ministry of Science and Higher Education (K.V.T. and N.N.S.).

The authors declare no conflict of interest.

## Author contributions

N.N.S. conceived the idea and supervised the study. N.N.S. and K.V.T. cloned and purified proteins, designed and performed most experiments. J.T. crystallized proteins. D.V.S. and K.V.T. performed and analyzed SPR experiments. E.G.M and K.V.T. performed and analyzed time-resolved fluorescence anisotropy measurements. N.N.S. solved and refined structures with input from A.A.A. N.N.S., E.G.M. and A.A.A interpreted all the results. N.N.S. wrote paper with input from A.A.A.

