## Supplementary figures and tables for "Molecular basis for the recognition of steroidogenic acute regulatory protein by the 14-3-3 protein family"

**SUPPLEMENTARY MATERIAL**

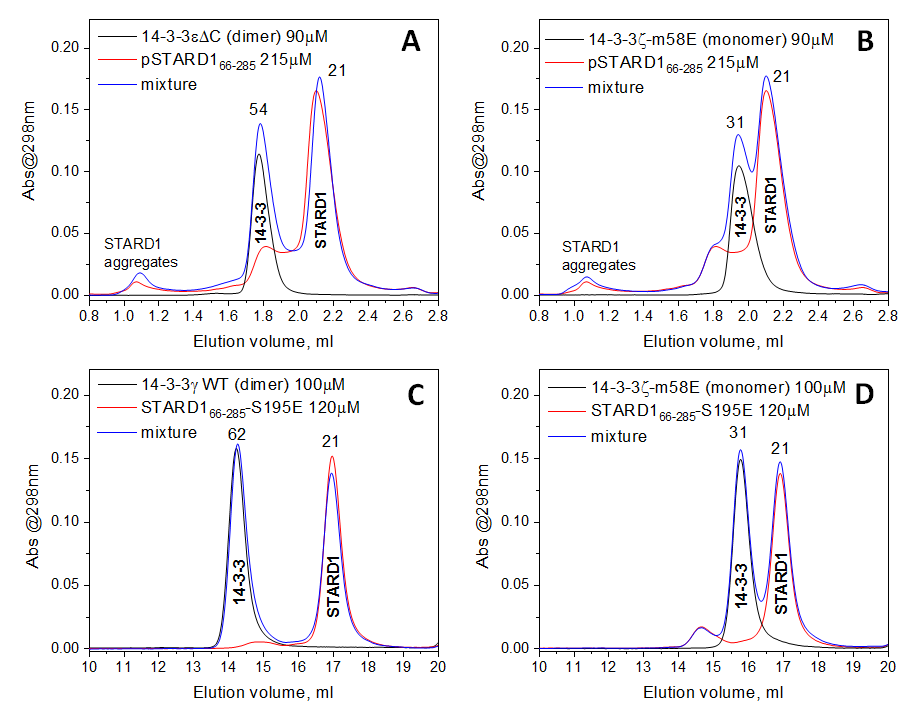

**Supplementary Fig. 1**. Analysis of STARD1_66-285_ interaction with 14-3-3. Size-exclusion chromatography profiles are shown for phosphorylated STARD1_66-285_ (A and B) or its phosphomimicking mutant S195E (C and D) with either dimeric (A and C) or monomeric 14-3-3 (B and D). An individual 14-3-3 variant, an individual STARD1 variant or their mixture (molar concentrations are indicated) were pre-incubated for 20 min at room temperature and then loaded on either a Superdex 200 Increase 5/150 column at 0.1 ml/min (A and B) or a Superdex 200 Increase 10/300 column at 1.5 ml/min (C and D). Mw values for the main peaks are indicated in kDa as determined from calibration. In all cases, no stable complexes could be observed. Neither of the STARD1/14-3-3 combinations used, yielded detectable interaction despite the very high protein concentrations used.

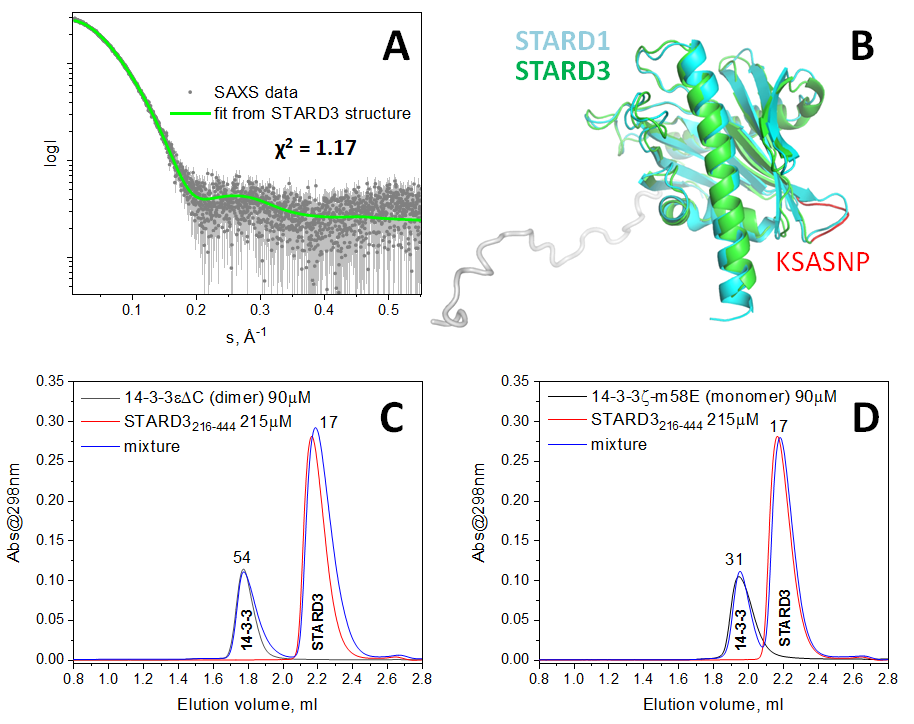

**Supplementary Fig. 2**. Analysis of STARD3_216-444_ interaction with 14-3-3. SAXS data for the STARD3 sample used in this study overlayed with a scattering plot calculated from crystal structure (PDB code 5I9J) using CORAL ^1^. B. Comparison of STARD3 and STARD1 (PDB 3P0L) structures (Cα RMSD ~1 Å) showing location of the 392-KSASNP-397 motif previously implicated in 14-3-3 binding ^2^. (C,D) SEC analysis of STARD3 interaction with the dimeric (C) or monomeric (D) 14-3-3. The indicated individual STARD3, 14-3-3 or their mixtures were pre-incubated for at least 20 min at room temperature and then loaded on a Superdex 200 Increase 5/150 column at a 0.1 ml/min flow rate. Mw values for the main peaks are indicated in kDa as derived from column calibration. Note the retardation of the STARD3 peak (apparent Mw of 17 kDa versus calculated Mw of 27.7 kDa), which is even more pronounced than previously reported for the human STARD1 ^3^. Neither of the STARD3/14-3-3 combinations used, yielded detectable interaction despite the very high protein concentrations used.

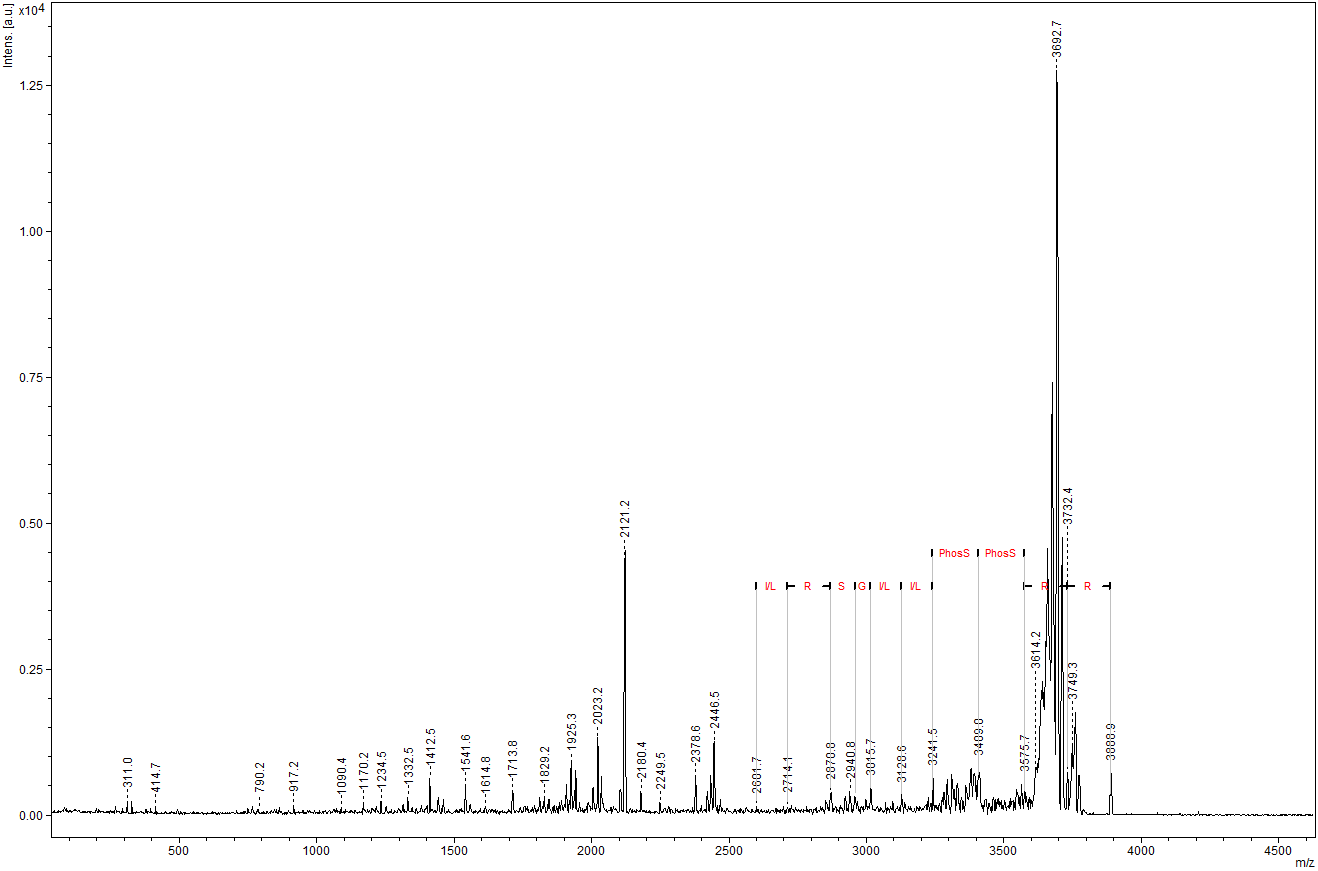

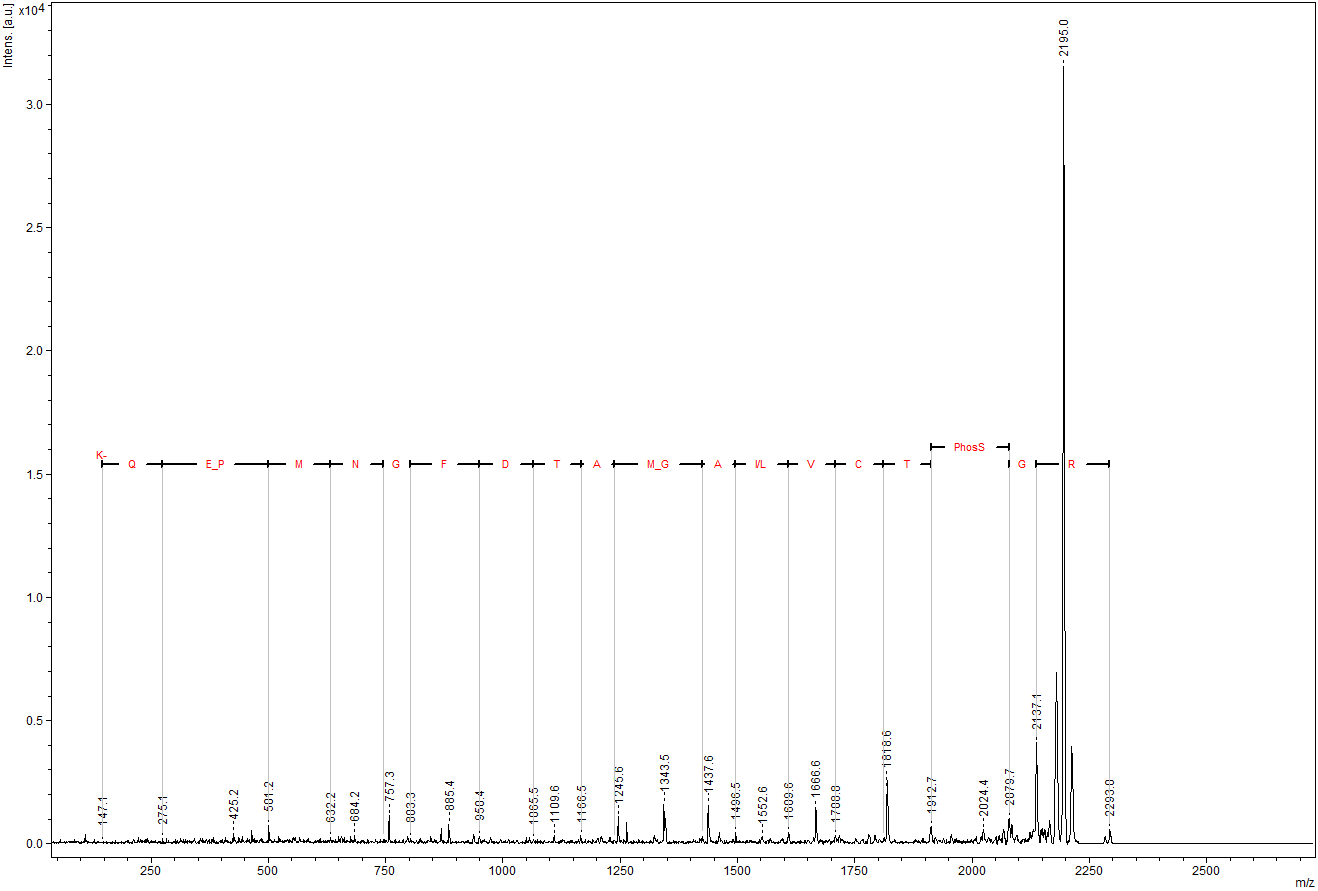

**Supplementary Fig 3**. Exemplary MS/MS spectra showing fragmentation of the doubly phosphorylated 3888.9 Da peptide containing pSer56 and pSer57 and of the singly phosphorylated 2293.0 Da peptide containing pSer195.

**Supplementary Table 1. Phosphopeptides of STARD1 verified by tandem mass-spectrometry after STARD1 in-gel trypsinolysis.**

| **Phosphorylated residues** | **Peptide** | **peptide mass, Da** | **Sample where observed** |
| --- | --- | --- | --- |
| Ser195 | RG**pS^195^**TCVLAGMATDFGNMPEQK RRG**pS^195^**TCVLAGMATDFGNMPEQK | 2293.0  2449.1 | in all STARD1 samples phosphorylated by PKA |
| Ser56+Ser57 | RR**pS^56^pS^57^**LLGSRLEETLYSDQELAYLQQGEEAMQK | 3888.9 | in STARD1_53-285_ |
| Ser57 | RG**pS^57^**LLGSR  RRG**pS^57^**LLGSR | 925.4  1081.5 | in STARD1 with S56G mutation* |

*-given this result, the phosphorylation of STARD1_46-285_-S56A was not additionally analyzed by mass-spectrometry.

**Supplementary Table 2. Crystallization conditions.**

| Designation | Chimera with the STARD1-Ser57 phosphopeptide | Chimera with the STARD1-Ser195 phosphopeptide |
| --- | --- | --- |
| Crystallization solution (resevoir) | 0.09 M NPS mix (0.3 M sodium nitrate, 0.3 M sodium phosphate dibasic, 0.3 M ammonium sulfate), 0.1 M Buffer mix (1.0 M Imidazole; MES), 25% v/v MPD; 25% polyethylene glycol 1000; 25% w/v polyethylene glycol 3350, pH 6.5 | 20% w/v polyethylene glycol 3350 |
| Crystal Handling | No cryo-solution | No cryo-solution |
| Resolution (Å) | 2.04 | 2.63 |
| Protein conc. (mg/mL) | 13.1 | 21.3 |
| Temperature (°C) | 6 | 6 |
| Growth Time (days) | 36 | 20 |

**Supplementary Table 3.** **X-ray data collection and refinement statistics.**

|  | Chimera with the STARD1-Ser57 phosphopeptide | Chimera with the STARD1-Ser195 phosphopeptide |
| --- | --- | --- |
| **Data collection** |  |  |
| Space group | *C 1 2 1* | *P 1 2_1_ 1* |
| Cell dimensions: *a*, *b*, *c* (Å) | 90.17, 78.35, 76.06 | 77.16, 74.95, 104.38 |
| α, β, γ (°) | 90, 100.42, 90 | 90, 97.98, 90 |
| Resolution range (Å)* | **49.14 – 2.04** [49.14-4.39] (2.11-2.04) | **53.51 – 2.63**  [53.52-5.66] (2.72-2.63) |
| Wavelength (Å) | 0.91587 | 0.97950 |
| *R*_merge_** | 0.106  [0.047] (1.31) | 0.150  [0.059] (1.38) |
| *R*_meas_ | 0.122  [0.054] (1.50) | 0.170  [0.069] (1.59) |
| *<I* / σ> | 8.4 (1.1) | 5.4 (1.1) |
| *CC_1/2_* | 1.0 (0.55) | 0.99 (0.48) |
| Completeness (%) | 99.7 (99.8) | 99.9 (99.6) |
| Redundancy | 4.2 (4.1) | 4.0 (4.0) |
| Wilson B-factor (Å^2^) | 36.2 | 63.2 |
| **Refinement** |  |  |
| No. of reflections: total | 33161 | 35345 |
| ‘free’ set | 1632 | 1811 |
| *R*_work(_%) | 18.5 | 21.7 |
| *R*_free (_%) | 22.9 | 26.0 |
| No. of 2:2 complexes /asu | 1 | 2 |
| No. of non-H atoms: protein/ligands/solvent | 3729 / 55 / 345 | 7515 / 40 / 111 |
| R.m.s.d. bond lengths (Å)/ angles (°) | 0.010 / 1.0 | 0.010 / 1.0 |
| Ramachandran favoured/ outliers (%) | 98.3 / 0.0 | 96.6 / 0.1 |
| Molprobity score/Clash score | 1.0 / 0.26 | 2.0 / 4.8 |
| PDB code | **6T5H** | **6T5F** |

*Statistics for the lowest and highest resolution shells are indicated in square brackets and parentheses, respectively

**All statistics as defined in XSCALE ^4^.
